# Allele-specific miRNA-binding analysis identifies candidate target genes for breast cancer risk

**DOI:** 10.1101/777318

**Authors:** Ana Jacinta-Fernandes, Joana M. Xavier, Ramiro Magno, Joel G. Lage, Ana-Teresa Maia

## Abstract

Most breast cancer (BC) risk-associated variants (raSNPs) identified in genome-wide association studies (GWAS) are believed to cis-regulate the expression of genes. We hypothesise that cis-regulatory variants contributing to disease risk may be affecting miRNA genes and/or miRNA-binding. To test this we adapted two miRNA-binding prediction algorithms — TargetScan and miRanda — to perform allele-specific queries, and integrated differential allelic expression (DAE) and expression quantitative trait loci (eQTL) data, to query 150 genome-wide significant (*P* ≤ 5 × 10^−8^) raSNPs, plus proxies. We found that no raSNP mapped to a miRNA gene, suggesting that altered miRNA targeting is an unlikely mechanism involved in BC risk. Also, 11.5% (6 out of 52) raSNPs located in 3’UTRs of putative miRNA target genes were predicted to alter miRNA∷mRNA pair binding stability in five candidate target genes. Of these, we propose *RNF115*, at locus 1q21.1, as a strong novel target gene associated with BC risk, and re-inforce the role of miRNA mediated cis-regulation at locus 19p13.11. We believe that integrating allele-specific querying in miRNA-binding prediction, and data supporting cis-regulation of expression, improves the identification of candidate target genes in BC risk, as well as in other common cancers and complex diseases.

## Introduction

In the last 10 years, genome-wide association studies (GWAS) identified hundreds of common low-penetrance variants to be associated with breast cancer (BC) risk^1^. Most of these risk-associated single-nucleotide polymorphisms (raSNPs) are located in non-coding regions^2^, often with no established, or easily perceived, biological function. Rather than altering protein sequence, and consequently protein function or structure, it seems that most raSNPs, or those in linkage disequilibrium with them, may act in cis to regulate the expression levels of target genes located distally and proximally^3–5^. The biological effect of raSNPs has so far been detected by expression quantitative trait loci analysis (eQTL)^3, 6–8^, but also, though less frequently, through the analysis of differential allelic expression (DAE)^9, 10^. A few functional studies for BC raSNPs have confirmed this cis-regulatory role, but have mainly focused on their potential to alter transcription factor binding sites^3, 6–8^. Nevertheless, genetic variation can modulate gene expression by several other mechanisms, such as microRNA-mediated regulation.

miRNAs are small non-coding RNA (ncRNA) molecules that bind messenger RNA (mRNA) complementary sequences and generally direct post-transcriptional silencing in the 3’ untranslated region (UTR) of target genes^11^. There is strong, albeit episodic, evidence of SNPs within miRNA genes and mRNA binding sites affecting the susceptibility to some cancers^12, 13^, including BC^14–16^. However, hitherto, the systematic analysis of BC risk loci via miRNA regulation is still lacking.

Here, we set out to evaluate the effect of common genetic variants associated with BC susceptibility on miRNA-regulatory mechanisms. Our initial list of raSNPs was established by selecting the 150 most significant (*P* ≤ 5 × 10^−8^) BC raSNPs from published GWAS (retrieved on 13/02/2017), along with their proxies in high LD. Next, we filtered these by genomic location, keeping those in or near miRNA genes and/or in protein-coding genes (PCGs; potential miRNA target sequences). Finally, we modified existing prediction tools to perform allele-specific miRNA target-prediction analysis. We used both miRNAs and putative mRNA target genes, expressed in normal breast tissue, and also cis-regulated genes as supported by DAE and eQTL data in normal breast tissue.

To our knowledge, this is the first systematic miRNA pathway-based study from published BC GWAS, using an allele-differential prediction analysis that is improved by the integration of DAE and eQTL data from normal breast tissue.

## Results

### Some BC risk variants locate to the 3’UTR of PCGs, but none to miRNA genes

To evaluate the contribution to BC risk of genetic variation modelling miRNA∷mRNA binding, we first assessed how many GWAS SNPs and their proxies were located in either miRNA genes or 3’UTRs of PCGs. We identified 2749 raSNPs, resulting from 150 BC GWAS-SNPs (Additional file 1: Table S1) and their proxies, of which almost one third (805 raSNPs) were solely annotated to “gene deserts” (585 raSNPs) or intergenic regions (220 raSNPs). The remainder 1944 raSNPs were located in either ncRNA genes or PCG’s (Figure 1), in a total of 161 unique Ensembl gene IDs, correspondent to 129 HGNC (HUGO Gene Nomenclature Committee) symbols.

**Figure 1.**
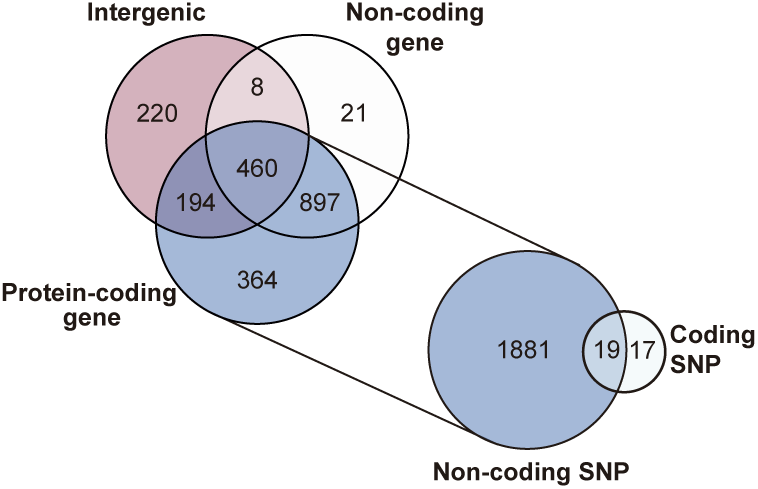
Genomic distribution of breast cancer risk-associated SNPs.

Next we assessed how many would change the miRNA gene sequence, thus affecting their biogenesis or target genes. Interestingly, none of the raSNPs mapped to miRNA genes, even after the LD threshold was lowered to *r*^2^ ≥ 0.2 when defining proxy SNPs (results not shown). This suggests that altered miRNA biogenesis or altered seed region sequence are unlikely mechanisms associated with BC risk. However, 13 SNPs were annotated as downstream or upstream variants of miRNA genes (Additional file 1: Table S2), raising the possibility of them being regulating the expression of the miRNA itself. However, we did not pursue this hypothesis further due to unavailability of DAE or eQTL data for these particular miRNA genes.

The vast majority of the raSNPs located within PCGs were in non-coding regions (1881 out of 1915, 98%) (Figure 1), consistent with previous reports^17^. SNPs located at the 3’UTR of the mRNA sequence of PCGs could potentially modify, create or destroy miRNA binding sites and we found 52 raSNPs (1.9% of total queried, 2.7% of total in PCG’s), at 16 risk loci, with at least one annotation at the 3’UTR of PCGs.

### Development and validation of allele-specific miRNA target-prediction analysis

raSNPs located at 3’UTR of PCGs were then evaluated for their potential to generate allelic differential miRNA-binding (Figure 2). To do so, we started by looking at existing miRNA-target prediction algorithms, but none could straightforwardly perform SNP allele queries in an automatic way (see Additional File 1: Table S3 for a systematic review). We, therefore, modified the input of two prediction algorithms, TargetScan^18^ and miRanda^19^, to account for SNP alleles queries and indirectly implemented them in R.

**Figure 2.**
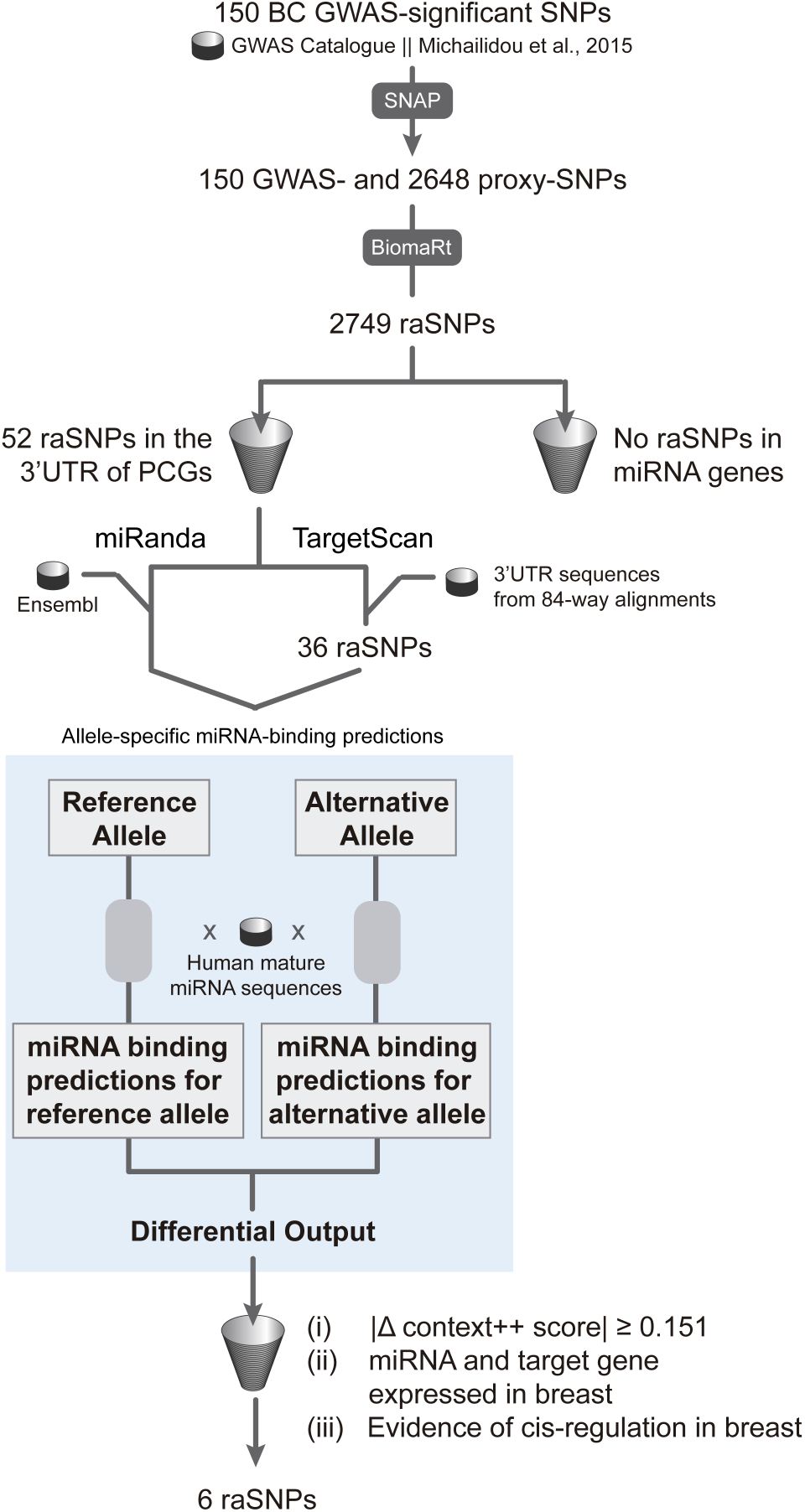
Schematic overview of the bioinformatics pipeline used for the prediction of allele-specific miRNA binding sites in breast cancer risk variants identified in GWAS. Genome-wide significant (*P* ≤ 5 × 10^−8^) SNPs associated with breast cancer risk in published GWAS were retrieved from the GWAS Catalog and from a GWAS meta-analysis^49^. Proxies in high linkage disequilibrium (*r*^2^ ≥ 0.8) were obtained through SNAP v2.2, using data from the pilot release of the 1000 Genomes Project for the CEU population. The biomaRt R package (v2.34.2) was used to retrieve genomic annotations from the Ensembl database v92. Risk-associated SNPs (raSNPs) were filtered for their location either in the 3’UTR of protein-coding genes (PCGs) or in miRNA genes. Next, allele-specific miRNA-binding predictions were performed by modifying the input of two prediction algorithms – TargetScan and miRanda. First, each raSNP allele (reference and alternative) located at the 3’UTR of PCGs was independently evaluated for putative miRNA-binding through the algorithms. Then, allele-specific miRNA binding predictions for each SNP were obtained by comparing each output file of corresponding SNP alleles and extracting their differences in miRNA binding. miRNA-binding predictions common to both algorithms were filtered for: (i) context++ score absolute difference (|Δ context++ score|) ≥ 0.151; (ii) miRNA expression (*RPM* > 1) in adjacent-normal breast tissue from the miRmine database and PCG expression (*TPM* > 1) in normal breast from the GTEx Project; and (iii) evidence of PCG cis-regulation in normal breast tissue from in-house differential allelic expression analysis and eQTL data from the GTEx Project.

To validate this approach of differential allelic miRNA binding querying, we ran the novel pipeline on seven SNPs that had previous functional validation supporting allele-specific miRNA binding. All seven SNPs were predicted to have allele-preferential binding of the corresponding miRNAs in at least one of the algorithms (Table 1), according to previous reports^13, 16, 20–24^. One of the SNPs previously published, and functionally validated, is rs11540855 in the *ABHD8* gene, which is in high LD (*r*^2^ = 0.86) with rs8170, a variant associated with risk to BC in *BRCA1* mutation carriers^25, 26^. Li et al.^20^ showed that the G allele of rs11540855 was preferentially bound by the hsa-miR-4707-3p agomir in the breast cancer cell line MCF116, leading to both decreased luciferase activity and ABHD8 protein levels. This is consistent with our prediction, using TargetScan, of hsa-miR-4707-3p specifically binding to the G allele of rs11540855 (Context++ Score = −0.225 for the G allele and no predicted binding for the allele A). The miRanda algorithm only predicted a minor difference in the maximum absolute minimum free energy (MFE) (−25.95 kcal/mol for the G allele and −23.93 kcal/mol for the A allele), although corroborating the preferential binding to the G allele (Table 1).

**Table 1.**
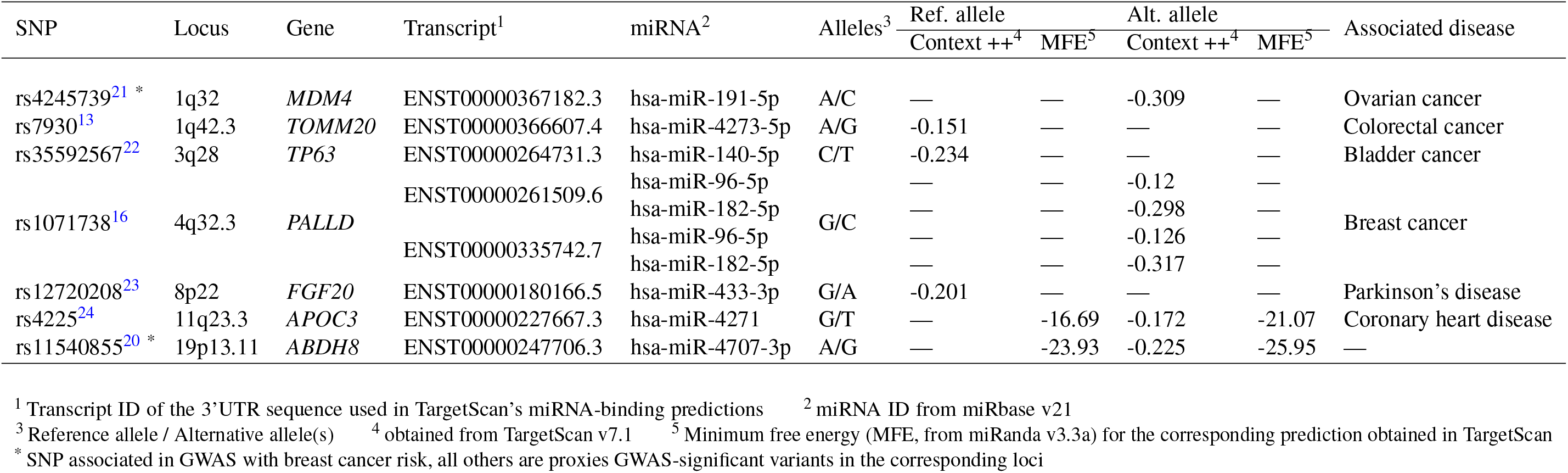
Previously functionally validated SNPs affecting miRNA binding evaluated by TargetScan and miRanda.

Another previously published prediction was that of the preferential binding of the hsa-miR-191-5p to the alternative allele (C allele) of rs4245739, located in the 3’UTR of the *MDM4* gene, in ovarian cancer cell lines^21^. This result was also concordant with our differential allelic miRNA binding predictions (context++ score of −0.309 for allele C vs no binding for allele A; Table 1).

The results from the validation step, presented in Table 1, suggest that functional allelic differences are easier to identify using the TargetScan algorithm. Additionally, they provided a guide to establish a biological significance threshold for the prediction scores, which we set at the maximum (i.e. the weakest binding prediction) TargetScan Context++ Score detected of −0.151, obtained for rs7930 in *TOMM20*^13^. This was later used to set the list of variants with stronger potential to affect miRNA binding.

### 5% of the tested raSNPs are predicted to alter miRNA binding

To assess how many of the 52 raSNPs located in the 3’UTR of PCGs (Additional file 1: Table S4) were likely to alter the miRNA∷mRNA pairing stability, the analysis pipeline was applied and the difference of scores obtained for each pair of alleles was calculated. Sixteen of these raSNPs could not be analysed by TargetScan, as they were 3’UTR variants of nonsense-mediated decay transcripts, which are excluded by this tool (Additional file 1: Table S4). Of the 36 raSNPs analysed for differential miRNA-binding by TargetScan, allele-specific context++ scores for a total of 311 different miRNAs (average nine miRNAs per SNP) were generated (Additional file 2). As for miRanda analysis, all 52 raSNPs generated maximum absolute MFE differences for a total of 2227 unique miRNAs (average 43 miRNAs per SNP; Additional file 3). Together, both algorithms commonly predicted a total of 160 combinations of gene-SNP-allele-miRNA. These were then filtered for the established TargetScan Context++ Score threshold, and evidence of both putative target mRNA (GTEx Project) and miRNA expression (miRmine database) in normal breast tissue, resulting in ten common predictions for seven raSNPs at six PCGs.

To identify candidate regulatory SNPs (rSNPs), we further filtered the resulting seven raSNPs based on previous evidence of cis-regulation of the target gene in breast tissue. For this purpose, we used the requirement of DAE of the target gene in normal breast tissue^27, 28^. Furthermore, we used normal breast tissue eQTL data^29^ as supporting evidence of cis-regulation. Overall, six BC raSNPs located in the 3’UTRs of five PCGs were predicted to modify miRNA∷mRNA pair binding stability in an allele-specific manner, with supporting evidence of cis-regulation of the putative target gene. These raSNPs correspond to five initial GWAS-significant associations in four BC risk loci (5% of the initial 83 BC GWAS loci) (Table 2). These variants were rs17354678 (in *RNF115*, at 1q21.1 locus), rs1019806 and rs6884232 (in *ATG10*, at 5q14.1-2 locus), rs3734805 (in *CCDC170*, at 6q25.1 locus), rs4808616 (in *ABHD8*, at 19p13.11 locus) and rs2385088 (in *ISYNA1*, at 19p13.11 locus).

**Table 2.**
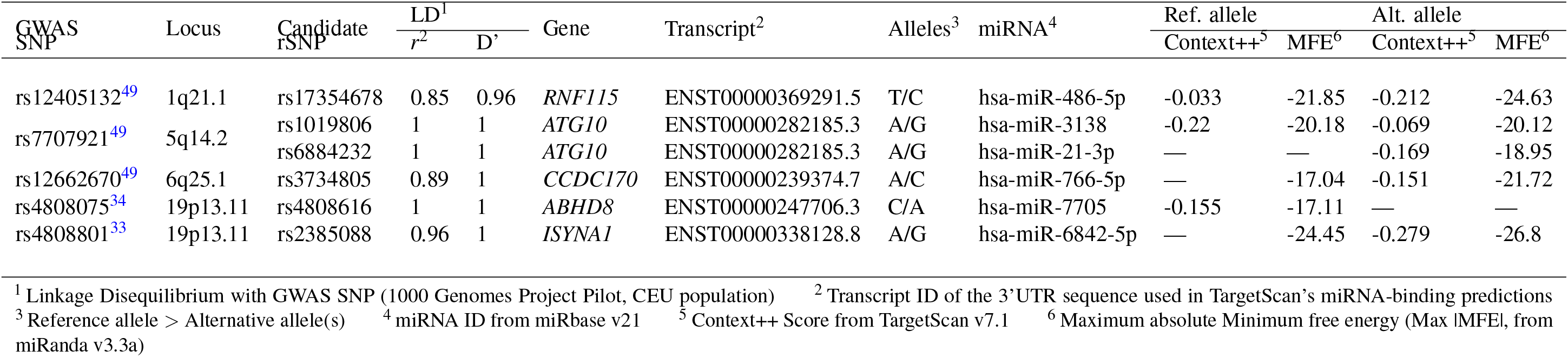
BC risk loci with putative allele-specific miRNA binding. Common miRNA-binding predictions found using TargetScan and miRanda algorithms. Results are filtered for (i) context++ score absolute fold change ≥ 0.151 between matching SNP alleles, (ii) miRNA expression with*RPM* > 1, using the miRmine dataset (SRX513286), and (iii) gene expression with *TPM* > 1 from the GTEx Project. Only genes with evidence of cis-regulation are shown. Results are ranked for decreasing context++ score.

### *RNF115* is a novel strong candidate target gene for BC risk

Following the canonical mechanism of action of miRNAs, we based our next analysis on the premise that the allele with preferential binding prediction would be the least expressed. Of the five PCGs with candidate rSNPs in their 3’UTRs with prediction for allelic-differential miRNA∷mRNA pair binding, the predictions for rs1019806 and rs6884232 (in *ATG10*), rs3734805 (in *CCDC170*) and rs2385088 (in *ISYNA1*), albeit supported by DAE evidence for the corresponding genes, were not in direct concordance with the preferential allelic expression pattern observed in normal breast tissue.

For *CCDC170*, the proposed rSNP rs3734805 is in very weak LD (*r*^2^≤0.2) with all the DAE variants analysed (Figure S2), and it is not an eQTL for the expression of any gene according to GTEx. Therefore, we could not establish direct association between the rSNP alleles and preferential allelic expression.

As for the two candidate rSNPs in *ATG10* (5q14.1-2 locus), both are reported eQTLs for *ATG10* expression using GTEx dataset (data not shown), with the alternative alleles associated with lower expression. This is concordant with the DAE data for rs1428940 (Figure S2), in high LD with these variants (*r*^2^ = 0.92 and 0.93, respectively). However, the predictions for allelic-preferential binding of miRNAs at rs6884232 is discordant with this evidence, as it is the reference allele which is predicted to have preferential binding. As for rs1019806, only the TargetScan prediction points to a concordant allelic difference in binding, whilst miRanda predicts almost no allelic difference.

For the pair hsa-miR-6842-5p∷*ISYNA1*, the G allele of rs2385088 was predicted to bind preferentially, but this was also the preferentially expressed allele in normal breast (Figure S2). Moreover, the DAE data was concordant with the role of *ISYNA1* as a reported tumour suppressor gene^30–32^, with protective G allele of the GWAS-significant variant rs4808801 (Odds Ratio (OR) for G allele = 0.93, 95% CI = [0.91-0.95] *P* = 5 × 10^−15^)^33^ in high LD with the preferentially expressed G allele of rs2385088 (*r*^2^ = 0.97, being G the alternative allele for both variants).

Regarding locus 19p13.11, a novel variant in the 3’UTR of *ABHD8*’s only expressed transcript ENST00000247706.3 (Figure S1), rs4808616, was predicted to have allelic-specific binding of hsa-miR-7705 to the reference C allele. The DAE measured at this variant indicates this allele is the less expressed (Figure 3A). rs4808616 is in complete LD with rs4808075, associated with cancer pleiotropy (OR for alternative C allele = 1.1, *P* = 4 × 10^−7^)^34^, suggesting that risk may be caused via increasing expression of *ABHD8*.

**Figure 3.**
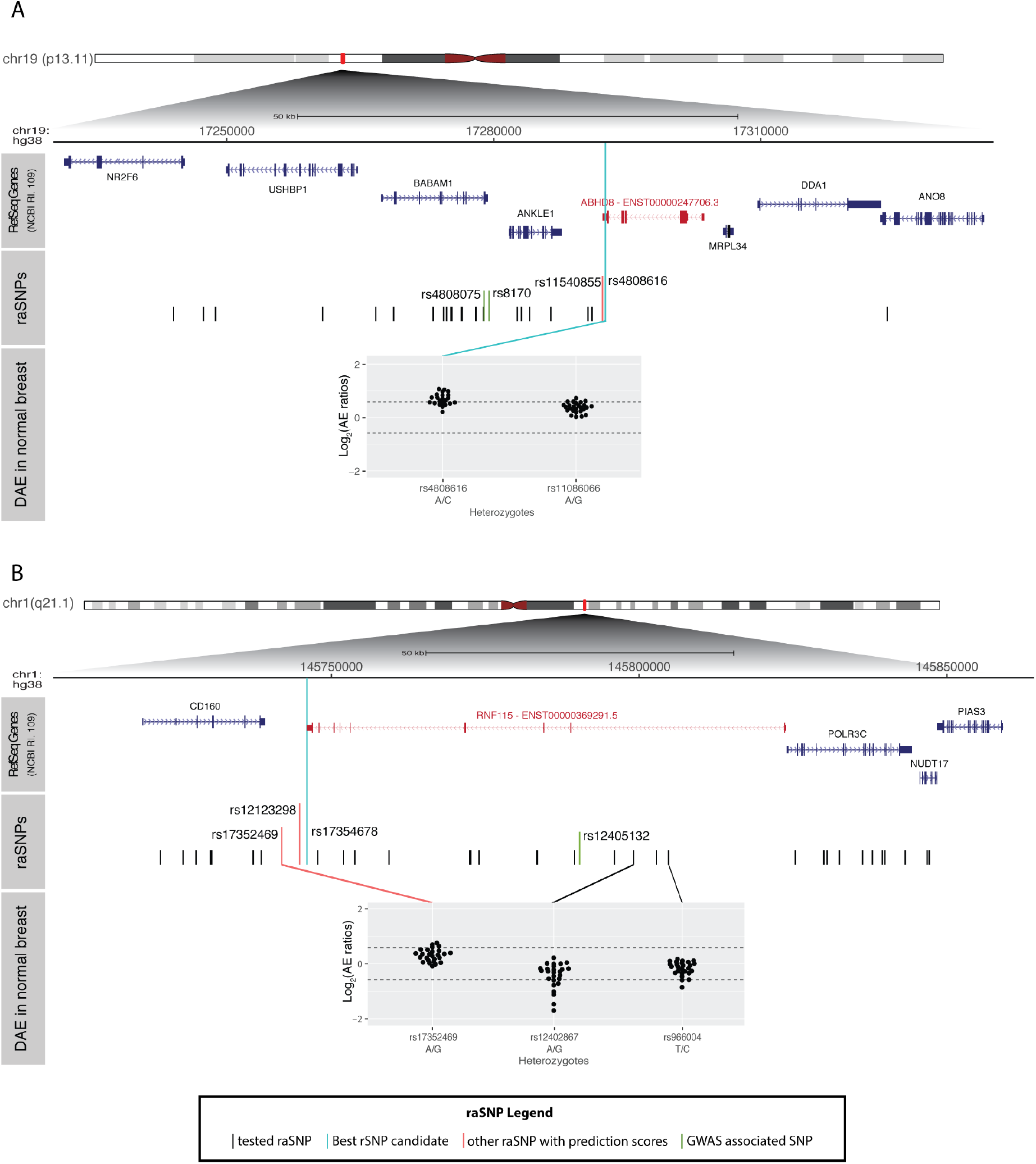
Breast cancer risk loci with strong predictions for allelic-differential binding of miRNA. **A** - BC risk locus 19p13.11 with candidate target gene *ABHD8* and **B** - BC risk locus 1q21.1 with candidate target gene *RNF115*. Both figures display a top panel with the genomic organisation of RefSeq genes (NCBI Release 109), with the candidate target genes in red and displaying the transcript predicted to have differential binding of miRNAs. The middle panel displays the genomic organisation of the tested raSNPs (GWAS-significant variants in green, plus proxy SNPs in high LD in black) in each locus, identified and tested in this study. raSNPs located in the 3’UTR of candidate target genes with the strongest predictions for allelic-differential binding of miRNAs (rSNP candidates) are indicated in cyan. raSNPs in red have weaker predictions for allelic-differential binding of miRNAs. The third panel shows the DAE data for heterozygous individuals tested for the SNP indicated immediately below. The alleles are indicated for each SNP in the order of the AE ratio calculated (i.e. A/G corresponds to the ratio of allele A by allele G). Dashed horizontal lines indicate the threshold for DAE set at 1.5 fold difference between alleles (*| log*_2_*AEratio*| = 0.58).

The last candidate rSNP, rs17354678, locates to the 3’UTR of *RNF115* and was predicted to have more stable pair binding of hsa-miR-486-5p∷*RNF115* in the presence of the alternative C allele, for the only protein coding transcript ENST00000369291.5 (Figure S1). rs17354678 is in high LD with variants for which DAE was detected, namely rs12402867 (*r*^2^ = 0.85) and rs17352469 (*r*^2^ = 0.9), both of which displayed preferential expression of the reference alleles (G and A, respectively) (Figure 3B). This was congruent with the prediction of preferential binding of the alternative C allele of rs17354678. This variant is also in high LD (*r*^2^ = 0.85) with the BC GWAS-significant SNP rs12405132 (OR for reference C allele = 1.03, 95% CI = [1.01-1.05], *P* = 6 × 10^−10^)^35^. Therefore, these results suggest that risk may be conferred by higher expression of *RNF115*.

## Discussion

GWAS have established the importance of normal non-coding genetic variation in common diseases, including BC. Currently, the challenges are to identify the true causal variant in risk-associated loci, as well as to determine the mechanism by which they act and the target genes they control. Our study focused on understanding the extent by which miRNA-mediated cis-regulation contributes to BC risk.

We found that none of the tested raSNPs from BC GWAS mapped to miRNA genes, suggesting that altered miRNA biogenesis, or targeting, are unlikely mechanisms to be involved in BC risk. However, this result could be due to the low representation of miRNA SNPs in the commercial genotyping arrays. On a biological perspective, however, miRNAs can bind hundreds of different mRNA targets^36^ across the genome, and variants in both the precursor elements, as well as the mature miRNA sequence, may drive changes in transcription in a much more widespread and significant manner, than those on target sequences^37^. Thus, we can hypothesise that SNPs in miRNA genes would have larger effect-sizes than those observed in GWAS, explaining their under-representation in these studies.

Next, we found thirteen BC raSNPs located upstream and downstream of miRNA genes, which could be regulating miRNA transcription, however most were also intronic variants of PCGs with evidence of cis-regulation from DAE data in normal breast tissue. This suggests that these raSNPs are more likely regulating PCG expression instead. Also, since the DAE data was generated using PCG exon-centric SNP microarrays, and as such did not cover the great majority of miRNA genes, we cannot exclude the possibility that our result is biased against miRNA gene cis-regulation. Further studies will be needed to evaluate whether raSNPs are located in regulatory regions affecting miRNA gene expression levels.

We also found 52 raSNPs located at the 3’UTR of PCGs, where miRNAs are generally known to bind^11^, supporting a possible role for miRNA mediated cis-regulation in BC predisposition. However, co-expression of both miRNAs and target sequences in the same tissue is imperative to validate such findings. While no comprehensive datasets are currently available to quantify the impact of miRNA expression on the putative target gene expression levels, we performed a qualitative filter for miRNA expression in adjacent-normal breast tissue^38^, as well as for the target-gene expression in breast mammary tissue^39^.

GWAS follow-up studies have mostly used eQTL mapping in normal tissue, to identify cis-acting variants and prioritize candidate cis-rSNPs for functional analysis^40–42^. However, cis-regulatory signals can be masked in eQTL studies by trans-acting factors or environmental effects^43^. Direct assessment of cis-regulation requires allele-specific approaches, such as DAE studies, where the effect of trans-regulation is eliminated when comparing the relative expression of two alleles in an heterozygous individual, within the same cellular context^44^. Here we combined both DAE and eQTL to filter our results, and we predict that six 3’UTR raSNPs have the potential to alter miRNA binding stability in five genes with evidence of cis-regulation in normal breast.

Our strongest result was obtained for raSNP rs17354678, mapping to the 3’UTR of the transcript ENST00000369291.5 of the *RNF115* gene, for which the reference allele was predicted to decrease the binding of hsa-miR-6842-5p in normal breast tissue. According to the DAE data, this allele is congruently associated with higher expression of *RNF115*, and is in high LD with the risk variant for BC, rs12405132^35^, supporting that risk might be conferred by upregulation of *RNF115*. *RNF115* encodes for the 3 ubiquitin ligase RING finger protein 115, which has been reported as upregulated in BC, particularly in estrogen receptor alpha positive tumours^45^. RNF115 has also been proposed to promote proliferation possibly through downregulation of the expression of the tumour suppressor p21^46^. Our data further supports RNF115’s role as an oncogene, as the predicted preferential binding of hsa-miR-486-5p and consequent lower expression of *RNF115*, is associated with protection against BC.

Additionally we also found a strong evidence that the reference allele of rs4808616, located at the 3’UTR of the transcript ENST00000247706.3 of the *ABHD8* gene could promote a binding site for hsa-miR-7705 in normal breast tissue. Expression and LD analysis with the BC risk variant support a role for miRNA mediated regulation and decreased expression of *ABHD8* in BC risk. This prediction is in accordance to what has been functionally validated for other candidate rSNP rs11540855^20^ in *ABHD8*. Previously, rs4808616 had been also functionally studied for mechanisms underlying pleiotropic risk to breast and ovarian cancer^47^. The authors found evidence of allelic expression and identified multiple risk alleles which they associated with increased *ABHD8* promoter activity. For rs4808616, in particular, the authors identified a link between the risk allele and higher expression of *ABHD8* through inclusion in a putative regulatory elements, but did not test for miRNA-mediated mechanisms. Nevertheless, all data, ours and from others, supports that increased expression of *ABHD8*, which encodes for a poorly studied lipase^48^, is linked to higher risk to BC. Furthermore, our study adds evidence for altered miRNA-binding through cis-regulatory variation as a mechanisms of risk in this locus.

In this study, we proposed a systematic post-GWAS framework focused on miRNA regulation, integrating DAE and eQTL data from normal tissue, to prioritize candidate rSNPs in already known risk loci. Although searching for altered transcription factor binding has been a popular approach following GWASes, other mechanisms, or even more than one at the same time, may be at play at susceptibility loci. Thus, it is important to look at the whole cis-regulation context when searching for the causal rSNP(s). Here, we showed that five genes, at four BC risk loci, have putative altered miRNA binding and that these genes have evidence of cis-regulation in normal breast tissue, supporting a functional role. In the future, it will be important to validate such findings through in vitro and in vivo assays.

Finally, our study provides a quick, powerful and systematic way of assessing the allelic-differential miRNA-mediated cis-regulation. As other common cancer GWASes have similar genomic distribution of risk variants to BC, it will be interesting to determine whether similar findings of putative altered miRNA regulation is also present in other common cancers.

## Methods

### BC risk loci dataset

GWAS-significant raSNPs for BC were retrieved from the NHGRI-EBI Catalog of published GWAS^2^, available at www.ebi.ac.uk/gwas (accessed on 13/02/2017), using *P* ≤ 5 × 10^−8^ and the catalogue traits “Breast cancer”, “Breast cancer (male)”, “Breast cancer (early onset)”, “Breast Cancer in *BRCA1* mutation carriers”, “Breast cancer in *BRCA2* mutation carriers”, “Breast cancer (estrogen-receptor negative, progesterone-receptor negative, and human epidermal growth factor-receptor negative)”, “Cancer” and “Cancer (pleiotropy)”. Furthermore, we retrieved the 15 SNPs previously identified by Michailidou and colleagues in a BC GWAS meta-analysis^49^, as well as those mentioned in their supplementary table 3, with a *P* ≤ 5 × 10^−8^, which included variants found associated by previous candidate gene association studies.

### Proxy SNP query

Proxy SNPs were identified using the SNP Annotation and Proxy Search^50^ online tool (version 2.2), available at archive. broadinstitute.org/mpg/snap/ldsearch.php, using genotype data from the pilot release of the 1000 Genomes Project^51^ for the CEU population (Utah residents with Northern and Western European ancestry), with a distance limit of 500kb and a LD threshold of *r*^2^ ≥ 0.8.

### Retrieval of SNP annotations

We used the getBM function from the biomaRt R package v2.34.2^52^ to retrieve the genomic annotations, alleles and variation consequence (Ensembl release 92^53^) of each SNP, as well as its molecular position (according to Ensembl release 75^54^). SNPs flagged by Ensembl^53, 55^ for containing errors or inconsistencies in their annotation were automatically excluded from further analysis.

### Allele-specific miRNA-binding predictions

raSNPs were filtered for the Sequence Ontology term “3_prime_UTR_variant” as a variant consequence^56^. For each allele of 3’UTR-located SNPs, miRNA∷mRNA interactions were searched using the default settings of the predictive algorithm TargetScan (Release 7.1)^18^ and custom settings for the miRanda (v3.3a) algorithm^19, 57^, as described below. The R code used to perform both analyses is available at https://github.com/maialab/postgwas-miRNA.

TargetScan predicts biological targets of miRNAs by searching for the presence of conserved canonical sites in 3’UTRs that match the “seed” region (2-7 nucleotides of the mature miRNA) of each miRNA^18^. The matches are made to human 3’UTRs from Gencode v19 (Ensembl 75) and their orthologs, as defined by UCSC whole-genome alignments (hg19)^18^. For each site, a context++ score is calculated; the lower the score, the higher the probability of effective target repression^18^. TargetScan source code, and accompanying datasets, were downloaded from http://www.targetscan.org/cgi-bin/targetscan/data_download.vert71.cgi, and run over the two alleles of each raSNP. Briefly, for each SNP located within a specific human 3’UTR multiple sequence alignment (as provided by TargetScan), independent text files containing either the reference or the alternative allele were generated according to source code instructions for the reference allele. We excluded from further analysis SNPs annotated as 3’UTR variants but not located within the available 3’UTR sequence alignments (Supplementary file 1: Table S4). Default instructions available for Context++ Score calculation were followed. For each miRNA-binding prediction, Context++ Score differences between correspondent SNP alleles were calculated.

miRanda detects potential miRNA target sites in genomic sequences by carrying a dynamic programming local alignment between query miRNA sequences and target mRNA sequences^19^. For each detected complementary match between a miRNA and a potential target gene two measures are calculated: (i) a score S based on sequence complementary, and (ii) the minimum free energy of the optimal miRNA-mRNA interaction. High S and low MFE values indicate potential target sites^19^. We used the getBM function of the R package biomaRt^52^ to retrieve a target-sequence centred on each allele and flanked by 25 nucleotides on either side, based on annotation in the Ensembl release 92^53^. miRNA mature sequences were retrieved from miRbase database (release 21, ftp://mirbase.org/pub/mirbase/21/) and filtered for Human^58^. miRanda’s software (v3.3a) was obtained from MicroRNA.org, a comprehensive resource of miRNA target predictions and expression profiles^59^, at www.microrna.org. A cut-off of S ≥ 80 and a MFE ≤ −16 kcal/mol was used as previously described^14^ to select for miRNA-binding. Maximum absolute MFE differences between matching SNP alleles for each SNP∷miRNA pair were calculated.

### miRNA and miRNA target gene expression

miRNA expression data, previously generated by miRNA-sequencing from pooled adjacent-normal breast tissue samples from eight BC patients^60^, was obtained from the miRmine Database^38^, available under the Sequence Read Archive ID SRX513286 (http://guanlab.ccmb.med.umich.edu/mirmine, accessed on 22/01/2017). miRNAs with expression values greater than 1 read per million were considered as expressed^61^.

Gene and transcript expression levels for 290 breast mammary tissue samples were obtained from the GTEx Portal (v7) at www.gtexportal.org on 12/12/2017^39^. Genes with median expression levels greater than 1 transcript per million were considered as expressed^62, 63^.

### Defining cis-regulation of gene expression

Cis-regulatory SNPs (rSNPs) act on regulatory elements, including promoters, enhancers and miRNA-binding sites, by modifying the binding affinity of trans-acting factors and thus specifically affecting gene expression in an allelic manner. This gives rise to unequal expression of transcribed alleles of the gene, a common feature in the human genome^64, 65^. Comparison of the relative expression of the two alleles in a heterozygous individual by DAE analysis is therefore a direct indicator of rSNPs acting in cis. Furthermore, measure of mRNA transcripts and association of their expression levels with genetic variants in eQTL studies can also indicate the presence of cis-rSNPs.

DAE analysis was performed at our laboratory^27^, with mRNA and DNA extracted from 64 normal breast tissue samples, using Illumina Exon510S-Duo SNP microarrays (data available from GEO at www.ncbi.nlm.nih.gov/geo/ under accession number GSE35023)^28^, and will be described elsewhere. The following equation was used for normalisation of allelic expression:

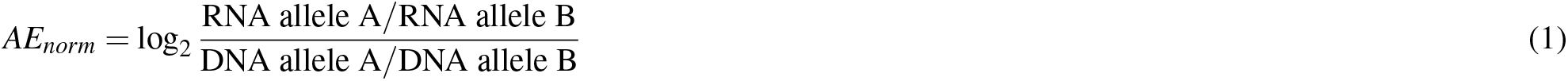

DAE was inferred when |*AE*_*norm*_| ≥ 0.58 (1.5 fold or greater allelic difference) for at least 10% of the heterozygotes and a minimum of 3 samples for a given SNP. Genes with at least one SNP with DAE were considered to show evidence of being cis-regulated.

cis-eQTL data (evaluated for ± 1Mb around the transcriptional start site of each gene) for 251 breast mammary tissue samples from GTEx’s v7 release was obtained from the GTEx Portal on 12/12/2017^29^. Genes whose expression levels were associated with at least one significant cis-eQTL, at a false discovery rate of ≤ 0.05, were selected.

## Acknowledgements

The authors would also like to thank Unidade de Apoio à Investigação (UAIC) at Universidade do Algarve (UAlg), in particular Mr Vitor Morais, and the Informatics Services of UAlg. We would also like to thank Dr Suet-Feung Chin and Dr Mae Goldgraben from the University of Cambridge for valuable discussions.

## Author contributions statement

AJ-F performed the computational work, and drafted the manuscript. JMX and RM performed the DAE analysis, critically revised the manuscript, and supervised the computational work. JGL conducted the computational analysis regarding SNPs near miRNA genes. A-TM conceived and directed the study, secured funding and drafted the manuscript. All authors read and approved the final manuscript.

## Additional information

### Availability of data and material

All datasets analysed in this study were obtained from publicly available databases and websites. SNPs associated with BC risk, as well as their proxies, were obtained from the GWAS Catalog website at www.ebi.ac.uk/gwas and from Broad Institute’s SNAP (v2.2) website at archive.broadinstitute.org/mpg/snap, respectively. SNP data was obtained from the Ensembl database (version 92 and 75) available at www.ensembl.org. The Pearl codes, as well as other datasets, used to perform TargetScan’s miRNA-binding predictions were obtained from the TargetScan (v7.1) website at www.targetscan.org/vert_71. miRanda’s software (v3.3a) was retrieved from MicroRNA.org at 34.236.212. 39/microrna/home.do. Mature miRNA sequences were retrieved from miRbase (release 21) at www.mirbase.org. The GTEx Project was supported by the Common Fund of the Office of the Director of the National Institutes of Health, and by NCI, NHGRI, NHLBI, NIDA, NIMH, and NINDS. Gene and transcript expression, and eQTL data for breast tissue from the GTEx Project (v7) were retrieved from the GTEx Portal at www.gtexportal.org. Files used to perform the analysis and generate this paper are available on GitHub at https://github.com/maialab/postgwas-miRNA. Further code may be made available upon request. Other detailed results are available in Additional files 1, 2 and 3.

## Additional Files

**Additional file 1 — Supplementary Tables and Figures (.pdf 1.1 MB)**

Tables S1-S4 and Figures S1-S2

**Additional file 2 (.txt 20 kb)**

A tab-separated text file with TargetScan’s miRNA binding predictions

**Additional file 3 (.txt 20 kb)**

A tab-separated text file with miRanda’s miRNA binding predictions

