## Supplementary File 1 for "Allele-specific miRNA-binding analysis identifies candidate target genes for breast cancer risk"

**Table S1.** 150 variants previously associated with breast cancer risk.

| SNP[1] | Locus | Position[2] | Allele[3] | AA[4] | MA[5] | Mapped Gene | Context | RAF[6] | *P* | OR [95% CI][7] | Trait[8] | Ref |
|---|---|---|---|---|---|---|---|---|---|---|---|---|
| rs186507655 | 1p36.33 | 1351675 | G/A | G | A | *DVL1 - MXRA8* | upstream gene variant | – | 5x10$^{-10}$ | – | Cancer (pleiotropy) | [1] |
| rs616488 | 1p36.22 | 10506158 | G/A | A | G | *PEX14* | intron variant | 0.665 | 1x10$^{-08}$ | 1.10 [1.06-1.14] | BC | [2] |
| | | | | | | | | 0.670 | 2x10$^{-10}$ | 1.06 [1.04-1.09] | BC | [3] |
| rs12118297 | 1p22.3 | 87313534 | T/G | G | T | *LOC101927844 - LMO4* | intergenic variant | 0.620 | 4x10$^{-08}$ | 1.10 [1.06-1.14] | BC | [4] |
| rs11552449 | 1p13.2 | 113905767 | C/G/T | C | T | *DCLRE1B* | missense variant | 0.170 | 2x10$^{-08}$ | 1.07 [1.04-1.09] | BC | [3] |
| rs11249433 | 1p11.2 | 121538815 | A/G | A | G | *EMBP1* | intron variant | 0.390 | 7x10$^{-10}$ | 1.16 [1.09-1.24] (Het) | BC | [5] |
| | | | | | | | | 0.400 | 2x10$^{-26}$ | 1.09 [1.07-1.11] | BC | [3] |
| rs12405132* | 1q21.1 | 145790097 | A/G | G | A | *RNF115* | intron variant | 0.640 | 8x10-$^{09}$ | 1.05 [1.03-1.08] | BC | [6] |
| rs12048493* | 1q21.2 | 149955122 | A/C | A | C | *OTUD7B* | intron variant | 0.340 | 1x10$^{-09}$ | 1.07 [1.05-1.10] | BC | [6] |
| rs6678914 | 1q32.1 | 202218048 | A/G | A | A | *LGR6* | intron variant | 0.590 | 1x10$^{-08}$ | 1.10 [1.06-1.13] | BC | [2] |
| rs4951011 | 1q32.1 | 203797203 | A/G | A | G | *ZC3H11A* | 5' UTR variant | 0.282 | 9x10$^{-09}$ | 1.09 [1.06-1.12] | BC | [7] |
| rs4245739 | 1q32.1 | 204549714 | A/C | A | C | *MDM4* | 3' UTR variant | 0.260 | 2x10$^{-12}$ | 1.14 [1.10-1.18] | BC | [2] |
| rs72755295* | 1q43 | 241870961 | A/G | A | G | *EXO1* | intron variant | 0.030 | 2x10$^{-08}$ | 1.15 [1.09-1.22] | BC | [6] |
| rs12710696 | 2p24.1 | 19121042 | C/T | C | T | *LOC105373455 - MIR4757* | intron variant | 0.360 | 5x10$^{-08}$ | 1.10 [1.06-1.13] | BC | [2] |
| rs4849887 | 2q14.2 | 120487546 | T/C | T | T | *LINC01101 - LOC105373585* | intergenic variant | 0.902 | 4x10$^{-11}$ | 1.10 [1.06-1.14] | BC | [3] |
| rs2016394 | 2q31.1 | 172108243 | A/G | G | A | *DLX2-AS1* | intron variant | 0.520 | 1x10$^{-08}$ | 1.05 [1.03-1.08] | BC | [3] |
| rs1550623 | 2q31.1 | 173348166 | G/A | A | G | *LOC100289479 - CDCA7* | intron variant | 0.840 | 3x10$^{-08}$ | 1.06 [1.03-1.09] | BC | [3] |
| rs13393577 | 2q34 | 212432139 | C/T | T | C | *ERBB4* | intron variant | 0.051 | 9x10$^{-14}$ | 1.53 [1.37-1.70] | BC | [8] |
| rs13387042 | 2q35 | 217041109 | G/A | G | A | *LOC105373874, LOC101928278* | intergenic variant | 0.500 | 1x10$^{-13}$ | 1.20 [1.14-1.26] | BC | [9] |
| | | | | | | | | 0.510 | 2x10$^{-08}$ | 1.25 [1.15-1.37] (Het) | BC | [10] |
| | | | | | | | | 0.520 | 2x10$^{-10}$ | 1.16 [1.11-1.22] | BC | [11] |
| | | | | | | | | 0.510 | 2x10$^{-57}$ | 1.14 [1.11-1.16] | BC | [3] |
| | | | | | | | | 0.490 | 2x10$^{-10}$ | 1.21 [1.14-1.29] | BC | [12] |
| rs16857609 | 2q35 | 217431785 | C/T | C | T | *DIRC3* | intron variant | 0.260 | 1x10$^{-15}$ | 1.08 [1.06-1.10] | BC | [3] |
| rs6762644 | 3p26.1 | 4700592 | A/G | A | G | *ITPR1* | intron variant | 0.400 | 2x10$^{-12}$ | 1.07 [1.04-1.09] | BC | [3] |
| rs481519 | 3p24.1 | 27285723 | C/T | T | C | *NEK10* | intron variant | – | 2x10$^{-09}$ | – | Cancer (pleiotropy) | [1] |
| rs653465 | 3p24.1 | 27302153 | C/T | T | C | *NEK10* | intron variant | 0.540 | 5x10$^{-12}$ | 1.18 [1.12-1.23] | BC (early onset) | [13] |
| rs4973768 | 3p24.1 | 27374522 | T/C | T | T | *SLC4A7* | 3' UTR variant | 0.490 | 2x10$^{-08}$ | 1.14 [1.09-1.19] | BC | [11] |
| | | | C/T | | | | | 0.470 | 2x10$^{-30}$ | 1.10 [1.08-1.12] | BC | [3] |
| rs12493607 | 3p24.1 | 30641447 | G/T/C | C | C | *TGFBR2* | intron variant | 0.350 | 2x10$^{-08}$ | 1.06 [1.03-1.08] | BC | [3] |
| rs6796502* | 3p21.31 | 46825376 | A/G | G | A | *PRSS43 - PRSS42* | downstream gene variant | 0.910 | 2x10$^{-08}$ | 1.09 [1.05-1.12] | BC | [6] |
| rs1053338* | 3p14.1 | 63982224 | A/G | A | G | *ATXN7* | missense variant | 0.130 | 9x10$^{-09}$ | 1.08 [1.05-1.11] | BC | [6] |
| rs7679673 | 4q24 | 105140377 | C/A | A | C | *LOC100288146 - TET2* | intron variant | – | 1x10$^{-10}$ | – | Cancer (pleiotropy) | [1] |
| rs9790517 | 4q24 | 105163621 | C/T | C | T | *TET2* | intron variant | 0.230 | 4x10$^{-08}$ | 1.05 [1.03-1.08] | BC | [3] |
| rs6828523 | 4q34.1 | 174925275 | A/C | C | A | *ADAM29* | intron variant | 0.870 | 4x10$^{-16}$ | 1.11 [1.09-1.15] | BC | [3] |
| rs10069690 | 5p15.33 | 1279675 | C/T | T | T | *TERT* | intron variant | 0.260 | 1x10$^{-10}$ | 1.18 [1.13-1.25] | BC | [14] |
| | | | | | | | | 0.321 | 5x10$^{-12}$ | 1.15 [1.11-1.20] | BC | [2] |
| | | | | | | | | 0.260 | 7x10$^{-09}$ | 1.06 [1.04-1.09] | BC | [3] |
| rs7725218 | 5p15.33 | 1282299 | G/A | G | A | *TERT* | intron variant | – | 2x10$^{-10}$ | – | Cancer (pleiotropy) | [1] |
| rs2736108* | 5p15.33 | 1297373 | T/C | C | T | *TERT - MIR4457* | upstream gene variant | 0.710 | 2x10$^{-09}$ | 1.06 [1.04-1.09] | BC | [6] |
| rs13162653* | 5p15.1 | 16187419 | A/T/G | G | T | *LOC401176 - NACAP6* | downstream gene variant | 0.550 | 1x10$^{-10}$ | 1.05 [1.03-1.08] | BC | [6] |
| rs2012709* | 5p13.3 | 32567626 | C/T | C | T | *SUB1* | intron variant | 0.460 | 6x10$^{-09}$ | 1.05 [1.03-1.08] | BC | [6] |
| rs4415084 | 5p12 | 44662413 | C/T | T | C | *LOC102723839 - RN7SL383P* | intergenic variant | 0.420 | 8x10$^{-11}$ | 1.17 [1.11-1.22] | BC | [11] |
| rs10941679 | 5p12 | 44706396 | A/G | A | G | *LOC102723839 - RN7SL383P* | intergenic variant | 0.250 | 2x10$^{-37}$ | 1.13 [1.10-1.15] | BC | [3] |
| rs7726159*† | 5p15.33 | 1282204 | C/A | C | A | TERT | intron variant | 0.340 | 3x10$^{-08}$ | 1.07 [1.02-1.11] | BC | [6,15] |
| rs16886034 | 5q11.2 | 56688029 | T/G/C | T | C | *LOC101928448 - LOC105378979* | regulatory region variant | 0.080 | 2x10$^{-09}$ | 1.36 [1.23-1.51] | BC (early onset) | [13] |
| rs16886113 | 5q11.2 | 56699208 | T/G | T | G | *LOC101928448 - LOC105378979* | intergenic variant | 0.080 | 4x10$^{-11}$ | 1.35 [1.23-1.47] | BC (early onset) | [13] |

| | | | | | | | | | | | | |
|---|---|---|---|---|---|---|---|---|---|---|---|---|
| rs16886181 | 5q11.2 | 56733416 | T/C | C | C | *LOC101928448 - LOC105378979* | intergenic variant | 0.180 | $9\times10^{-14}$ | 1.26 [1.18-1.34] | BC (early onset) | [13] |
| rs889312 | 5q11.2 | 56736057 | A/C | A | C | *LOC101928448 - LOC105378979* | regulatory region variant | 0.290 | $1\times10^{-08}$ | 1.29 [ – ] | BC (early onset) | [13] |
| | | | | | | | | 0.280 | $7\times10^{-20}$ | 1.13 [1.10-1.16] | BC | [16] |
| | | | | | | | | 0.280 | $5\times10^{-09}$ | 1.22 [1.14-1.30] | BC | [12] |
| | | | | | | | | 0.280 | $3\times10^{-36}$ | 1.12 [1.10-1.15] | BC | [3] |
| rs1862626 | 5q11.2 | 56737113 | G/T | T | G | *LOC101928448 - LOC105378979* | regulatory region variant | – | $4\times10^{-12}$ | – | Cancer (pleiotropy) | [1] |
| rs16886364 | 5q11.2 | 56826517 | A/G | A | G | *MAP3K1* | intron variant | 0.070 | $5\times10^{-12}$ | 1.36 [1.25-1.48] | BC (early onset) | [13] |
| rs16886397 | 5q11.2 | 56838449 | A/G | A | G | *MAP3K1* | intron variant | 0.070 | $4\times10^{-12}$ | 1.36 [1.25-1.49] | BC (early onset) | [13] |
| rs1017226 | 5q11.2 | 56857565 | T/C | T | C | *MAP3K1* | intron variant | 0.080 | $6\times10^{-11}$ | 1.33 [1.22-1.45] | BC (early onset) | [13] |
| rs2229882 | 5q11.2 | 56872885 | C/T | C | T | *MAP3K1* | synonymous variant | 0.060 | $1\times10^{-14}$ | 1.45 [1.32-1.60] | BC (early onset) | [13] |
| rs16886448 | 5q11.2 | 56874986 | C/G | C | G | *MAP3K1* | intron variant | 0.070 | $2\times10^{-12}$ | 1.37 [1.25-1.49] | BC (early onset) | [13] |
| rs3822625 | 5q11.2 | 56882284 | A/G | A | G | *MAP3K1* | synonymous variant | 0.070 | $5\times10^{-12}$ | 1.36 [1.24-1.48] | BC (early onset) | [13] |
| rs12655019 | 5q11.2 | 56899963 | A/G | A | G | *LOC105378980* | downstream gene variant | 0.100 | $3\times10^{-10}$ | 1.27 [1.18-1.37] | BC (early onset) | [13] |
| rs7726354 | 5q11.2 | 56960656 | C/T | C | T | *MIER3 - LOC100130001* | intron variant | 0.060 | $7\times10^{-11}$ | 1.37 [1.24-1.50] | BC (early onset) | [13] |
| rs10472076 | 5q11.2 | 58888234 | T/A/C | T | C | *RAB3C - PDE4D* | intergenic variant | 0.380 | $3\times10^{-08}$ | 1.05 [1.03-1.07] | BC | [3] |
| rs1353747 | 5q11.2 | 59041654 | G/T | T | G | *PDE4D* | intron variant | 0.905 | $3\times10^{-08}$ | 1.09 [1.05-1.12] | BC | [3] |
| rs7707921* | 5q14.2 | 82242227 | T/A | A | T | *ATG10* | intron variant | 0.770 | $5\times10^{-11}$ | 1.08 [1.05-1.10] | BC | [6] |
| rs10474352 | 5q14.3 | 91436408 | T/C | C | T | *ARRDC3-AS1 - RAB5CP2* | intron variant | 0.482 | $2\times10^{-09}$ | 1.09 [1.06-1.12] | BC | [7] |
| rs1432679 | 5q33.3 | 158817075 | T/C | C | T | *EBF1* | intron variant | 0.430 | $2\times10^{-14}$ | 1.07 [1.05-1.09] | BC | [3] |
| rs11242675 | 6p25.3 | 1318643 | C/T | C | C | *FOXQ1 - LINC01394* | downstream gene variant | 0.610 | $7\times10^{-09}$ | 1.06 [1.04-1.09] | BC | [3] |
| rs204247 | 6p23 | 13722291 | A/G | A | G | *RANBP9 - MCUR1* | intergenic variant | 0.430 | $8\times10^{-09}$ | 1.05 [1.03-1.07] | BC | [3] |
| rs9257408* | 6p22.1 | 28958443 | G/C | G | G | *TRM-CAT3-2 - KRT18P1* | regulatory region variant | 0.380 | $5\times10^{-08}$ | 1.05 [1.03-1.08] | BC | [6] |
| rs17529111* | 6q14.1 | 81418669 | T/C | T | C | *LOC648934 - LOC105377871* | intergenic variant | 0.220 | $2\times10^{-10}$ | 1.06 [1.04-1.08] | BC | [6] |
| rs17530068 | 6q14.1 | 81483392 | T/C | T | C | *LOC105377871* | intergenic variant | 0.220 | $8\times10^{-09}$ | 1.05 [1.03-1.08] | BC | [3] |
| rs2180341 | 6q22.33 | 127279485 | A/G | A | G | *RNF146* | intron variant | 0.210 | $3\times10^{-08}$ | 1.41 [1.25-1.59] | BC | [17] |
| rs9485372 | 6q25.1 | 149287738 | A/G | G | A | *TAB2* | intron variant | 0.550 | $4\times10^{-12}$ | 1.11 [1.09-1.15] | BC | [18] |
| rs3757318 | 6q25.1 | 151592978 | G/A | G | A | *CCDC170* | intron variant | 0.070 | $2\times10^{-21}$ | 1.16 [1.12-1.21] | BC | [3] |
| rs12662670* | 6q25.1 | 151597721 | T/C/G | T | G | *CCDC170* | intron variant | 0.070 | $7\times10^{-27}$ | 1.17 [1.13-1.22] | BC | [6] |
| rs2046210 | 6q25.1 | 151627231 | G/A | A | A | *CCDC170 - ESR1* | intergenic variant | 0.350 | $7\times10^{-15}$ | 1.22 [1.16-1.29] | BC | [4] |
| | | | | | | | | 0.423 | $5\times10^{-16}$ | 1.15 [1.11-1.19] | BC | [2] |
| | | | | | | | | 0.370 | $2\times10^{-15}$ | 1.29 [1.21-1.37] | BC | [19] |
| | | | A/G | | | | | 0.080 | $5\times10^{-09}$ | 1.28 [1.18-1.39] | BC in BRCA1 mutation carriers | [20] |
| rs140068132 | 6q25.1 | 151633699 | G/A | A | G | *CCDC170 - ESR1* | intergenic variant | 0.930 | $9\times10^{-18}$ | 1.67 [1.49-1.89] | BC | [21] |
| rs9383938 | 6q25.1 | 151666222 | G/T | G | T | *ESR1* | intron variant | – | $2\times10^{-10}$ | 1.28 [ – ] | BC | [22] |
| rs6964587*† | 7q21.2 | 92001306 | G/T | T | T | AKAP9 | missense variant | 0.390 | $9\times10^{-11}$ | 1.03 [1.02-1.05] | BC | [6,23] |
| rs4593472* | 7q32.3 | 130982362 | T/C | C | T | *LINC-PINT* | intron variant | 0.650 | $2\times10^{-09}$ | 1.05 [1.03-1.06] | BC | [6] |
| rs720475 | 7q35 | 144377836 | A/G | G | A | *ARHGEF5* | intron variant | 0.750 | $7\times10^{-11}$ | 1.06 [1.04-1.09] | BC | [3] |
| rs9693444 | 8p12 | 29652100 | C/A | C | A | *RPL17P33 - LINC00589* | intergenic variant | 0.320 | $9\times10^{-14}$ | 1.07 [1.05-1.09] | BC | [3] |
| rs13365225* | 8p11.23 | 37000965 | G/A | A | G | *RPL26P25 - LOC105379377* | intergenic variant | 0.830 | $1\times10^{-08}$ | 1.05 [1.02-1.08] | BC | [6] |
| rs6472903 | 8q21.13 | 75318066 | G/T | T | G | *HIGD1AP6 - PKMP4* | intron variant | 0.820 | $2\times10^{-17}$ | 1.10 [1.08-1.12] | BC | [3] |
| rs2943559 | 8q21.13 | 75505702 | A/G | A | G | *HNF4G* | intron variant | 0.070 | $6\times10^{-15}$ | 1.13 [1.09-1.17] | BC | [3] |
| rs13267382* | 8q23.3 | 116197325 | G/A | A | G | *LINC00536* | intron variant | 0.360 | $2\times10^{-08}$ | 1.05 [1.03-1.07] | BC | [6] |
| rs13281615 | 8q24.21 | 127343372 | A/G | A | G | *CASC21, CASC8* | intron variant | 0.400 | $5\times10^{-12}$ | 1.08 [1.05-1.11] | BC | [16] |
| | | | | | | | | 0.410 | $1\times10^{-27}$ | 1.09 [1.07-1.12] | BC | [3] |
| rs1562430 | 8q24.21 | 127375606 | C/T | T | C | *CASC21, CASC8* | intron variant | 0.600 | $3\times10^{-11}$ | 1.16 [1.11-1.22] | BC | [11] |
| rs2392780 | 8q24.21 | 127375779 | G/A | G | G | *CASC21, CASC8* | intron variant | 0.600 | $1\times10^{-08}$ | 1.15 [1.10-1.20] | BC (early onset) | [13] |
| rs11780156 | 8q24.21 | 128182395 | C/T | C | T | *MIR1208 - RN7SKP226* | regulatory region variant | 0.160 | $3\times10^{-11}$ | 1.07 [1.04-1.10] | BC | [3] |
| rs1011970 | 9p21.3 | 22062135 | G/T | G | T | *CDKN2B-AS1* | intron variant | 0.170 | $3\times10^{-08}$ | 1.09 [1.04-1.14] | BC | [12] |
| rs10759243 | 9q31.2 | 107543834 | C/T/A | A | A | *LOC105376206 - LOC105376205* | upstream gene variant | 0.390 | $1\times10^{-08}$ | 1.06 [1.03-1.08] | BC | [3] |
| rs865686 | 9q31.2 | 108126198 | G/A/T | T | G | *LOC105376214* | intergenic variant | 0.610 | $2\times10^{-10}$ | 1.12 [1.09-1.18] | BC | [11] |

| | | | | | | | | | | | | |
|---|---|---|---|---|---|---|---|---|---|---|---|---|
| | | | | | | | | 0.620 | 1x10$^{-34}$ | 1.12 [1.10-1.14] | BC | [3] |
| rs7072776 | 10p12.31 | 21744013 | G/A | A | A | *LOC107984214* | downstream gene variant | 0.290 | 4x10$^{-14}$ | 1.07 [1.05-1.09] | BC | [3] |
| rs11814448 | 10p12.31 | 22026914 | A/C | G | C | *DNAJC1 - ADIPOR1P1* | intergenic variant | 0.020 | 9x10$^{-16}$ | 1.26 [1.18-1.35] | BC | [3] |
| rs10822013 | 10q21.2 | 62492218 | C/T | C | T | *ZNF365* | intron variant | 0.470 | 6x10$^{-09}$ | 1.12 [1.06-1.18] | BC | [24] |
| rs10995190 | 10q21.2 | 62518923 | A/G | G | A | *ZNF365* | intron variant | 0.850 | 5x10$^{-15}$ | 1.16 [1.10-1.22] | BC | [12] |
| | | | | | | | | 0.840 | 1x10$^{-36}$ | 1.16 [1.14-1.19] | BC | [3] |
| rs704010 | 10q22.3 | 79081391 | C/T | C | T | *ZMIZ1* | intron variant | 0.390 | 4x10$^{-09}$ | 1.07 [1.03-1.11] | BC | [12] |
| | | | | | | | | 0.380 | 7x10$^{-22}$ | 1.08 [1.06-1.10] | BC | [3] |
| rs7904519 | 10q25.2 | 113014168 | A/G | G | G | *TCF7L2* | intron variant | 0.460 | 3x10$^{-08}$ | 1.06 [1.04-1.08] | BC | [3] |
| rs11199914 | 10q26.12 | 121334387 | T/C | T | T | *LOC105378523 - RN7SKP167* | intergenic variant | 0.680 | 2x10$^{-08}$ | 1.05 [1.03-1.08] | BC | [3] |
| rs11200014 | 10q26.13 | 121575416 | G/T/A | G | A | *FGFR2* | intron variant | — | 8x10$^{-35}$ | — | Cancer (pleiotropy) | [1] |
| rs2981579 | 10q26.13 | 121577821 | G/A | G | A | *FGFR2* | intron variant | 0.420 | 4x10$^{-31}$ | 1.43 [1.35-1.53] | BC | [12] |
| | | | | | | | | 0.400 | 2x10$^{-170}$ | 1.27 [1.24-1.29] | BC | [3] |
| | | | | | | | | 0.550 | 3x10$^{-11}$ | 1.19 [1.11-1.23] | BC | [4] |
| | | | | | | | | 0.410 | 2x10$^{-10}$ | 1.17 [1.07-1.27] (Het) | BC | [5] |
| rs2981578 | 10q26.13 | 121580797 | A/T/C | C | T | *FGFR2* | intron variant | 0.514 | 1x10$^{-12}$ | 1.23 [1.16-1.30] | BC | [25] |
| rs2981575 | 10q26.13 | 121586602 | G/A | G | G | *FGFR2* | intron variant | 0.420 | 1x10$^{-08}$ | 1.28 [1.18-1.39] | BC in BRCA2 mutation carriers | [26] |
| rs1219648 | 10q26.13 | 121586676 | A/G | A | G | *FGFR2* | intron variant | 0.420 | 2x10$^{-13}$ | 1.32 [1.22-1.42] | BC | [27] |
| | | | | | | | | 0.390 | 6x10$^{-09}$ | 1.17 [1.11-1.23] | BC | [4] |
| | | | | | | | | 0.400 | 1x10-$^{10}$ | 1.20 [1.07-1.42] | BC | [28] |
| | | | A/G | | | | | 0.420 | 1x10$^{-30}$ | 1.31 [1.25-1.37] | BC | [11] |
| rs2912774 | 10q26.13 | 121589148 | A/G/T | T | T | *FGFR2* | intron variant | 0.440 | 3x10$^{-27}$ | 1.29 [1.23-1.35] | BC (early onset) | [13] |
| rs2981582 | 10q26.13 | 121592803 | G/A | G | A | *FGFR2* | intron variant | 0.380 | 2x10$^{-76}$ | 1.26 [1.23-1.30] | BC | [16] |
| | | | | | | | | 0.680 | 2x10$^{-08}$ | 1.18 [1.11-1.23] | BC | [4] |
| rs3817198 | 11p15.5 | 1887776 | T/C | T | C | *LSP1* | intron variant | 0.300 | 3x10$^{-09}$ | 1.07 [1.04-1.11] | BC | [16] |
| | | | | | | | | 0.310 | 2x10$^{-11}$ | 1.07 [1.05-1.09] | BC | [3] |
| rs3903072 | 11q13.1 | 65815595 | T/G | G | T | *OVOL1 - SNX32* | regulatory region variant | 0.530 | 9x10$^{-12}$ | 1.05 [1.04-1.08] | BC | [3] |
| rs7931342 | 11q13.3 | 69227030 | T/G | G | T | *LOC105369366 - LOC105369367* | intergenic variant | — | 4x10$^{-14}$ | — | Cancer (pleiotropy) | [1] |
| rs537626 | 11q13.3 | 69492927 | G/C | G | C | *LINC01488* | intergenic variant | 0.180 | 2x10$^{-15}$ | 1.29 [1.21-1.37] | BC (early onset) | [13] |
| rs614367 | 11q13.3 | 69513996 | C/T | C | | *LINC01488 - CCND1* | intergenic variant | 0.150 | 2x10$^{-63}$ | 1.21 [1.18-1.24] | BC | [3] |
| | | | | | | | | 0.150 | 3x10$^{-15}$ | 1.15 [1.10-1.20] | BC | [12] |
| | | | | | | | | 0.160 | 1x10$^{-08}$ | 1.34 [ — ] | BC (early onset) | [13] |
| rs78540526* | 11q13.3 | 69516650 | C/T | C | T | *LINC01488 - CCND1* | regulatory region variant | 0.080 | 2x10$^{-86}$ | 1.34 [1.29-1.38] | BC | [6] |
| rs554219* | 11q13.3 | 69516874 | C/A/T/G | C | | *LINC01488 - CCND1* | regulatory region variant | 0.120 | 2x10$^{-81}$ | 1.26 [1.23-1.30] | BC | [6] |
| rs75915166* | 11q13.3 | 69564393 | C/A | C | A | *LINC01488 - CCND1* | intergenic variant | 0.060 | 1x10$^{-57}$ | 1.31 [1.26-1.36] | BC | [6] |
| rs148883465 | 11q22.3 | 103813371 | A/G | A | G | *LOC105369463 - LOC102723862* | intron variant | — | 3x10$^{-08}$ | — | Cancer (pleiotropy) | [1] |
| rs11820646 | 11q24.3 | 129591276 | T/C | T | T | *RPS27P20 - LINC01395* | upstream gene variant | 0.590 | 1x10$^{-09}$ | 1.05 [1.03-1.08] | BC | [3] |
| rs12422552 | 12p13.1 | 14260997 | G/C | C | C | *GNAI2P1 - RPL30P11* | regulatory region variant | 0.260 | 4x10$^{-08}$ | 1.05 [1.03-1.07] | BC | [3] |
| rs10771399 | 12p11.22 | 28002147 | A/G | A | G | *PTHLH - LOC105369710* | intergenic variant | | 2x10$^{-08}$ | 1.39 [1.25-1.56] | BC (ER-, PR, and HER-) | [29] |
| | | | G/A | | | | | 0.895 | 2x10$^{-12}$ | 1.20 [1.15-1.27] | BC | [2] |
| | | | | | | | | 0.880 | 8x10$^{-31}$ | 1.16 [1.14-1.20] | BC | [3] |
| rs73110464 | 12q13.13 | 52918828 | C/T | C | T | *KRT8* | intron variant | — | 2x10$^{-15}$ | — | Cancer (pleiotropy) | [1] |
| rs17356907 | 12q22 | 95633983 | G/A | A | G | *USP44 - PGAM1P5* | intron variant | 0.700 | 2x10$^{-22}$ | 1.10 [1.08-1.12] | BC | [3] |
| rs1292011 | 12q24.21 | 115398717 | G/A | G | G | *LOC105370003* | regulatory region variant | 0.580 | 9x10$^{-22}$ | 1.09 [1.06-1.11] | BC | [3] |
| rs56084662 | 13q13.1 | 32295727 | G/A | G | A | *FRY* | 3' UTR variant | — | 4x10$^{-09}$ | — | Cancer (pleiotropy) | [1] |
| rs11571818 | 13q13.1 | 32394673 | T/C | T | C | *BRCA2* | intron variant | — | 5x10$^{-10}$ | — | Cancer (pleiotropy) | [1] |
| rs11571833 | 13q13.1 | 32398489 | A/T | A | T | *BRCA2* | stop gained | 0.008 | 5x10$^{-08}$ | 1.26 [1.14-1.39] | BC | [3] |
| | | | | | | | | — | 8x10$^{-12}$ | 1.6 [ — ] | Cancer | [1] |
| rs2236007 | 14q13.3 | 36663564 | A/G | G | A | *PAX9* | intron variant | 0.790 | 2x10$^{-13}$ | 1.08 [1.05-1.10] | BC | [3] |
| rs2588809 | 14q24.1 | 68193711 | C/T | T | T | *RAD51B* | intron variant | 0.160 | 1x10$^{-10}$ | 1.08 [1.05-1.11] | BC | [3] |

| | | | | | | | | | | | | |
|---|---|---|---|---|---|---|---|---|---|---|---|---|
| rs1314913 | 14q24.1 | 68232877 | C/T | C | T | *RAD51B* | intron variant | — | $3x10^{-13}$ | 1.57 [1.39-1.77] | BC (male) | [30] |
| rs11844632 | 14q24.1 | 68559662 | G/A | A | A | *RAD51B* | intron variant | — | $3x10^{-10}$ | — | Cancer (pleiotropy) | [1] |
| rs999737 | 14q24.1 | 68567965 | T/C | C | T | *RAD51B* | intron variant | 0.770 | $3x10^{-19}$ | 1.09 [1.06-1.11] | BC | [3] |
| rs941764 | 14q32.11 | 91374725 | A/G | G | G | *CCDC88C* | intron variant | 0.340 | $4x10^{-10}$ | 1.06 [1.04-1.09] | BC | [3] |
| rs11627032* | 14q32.12 | 92637727 | C/T | T | C | *RIN3* | intron variant | 0.740 | $4x10^{-09}$ | 1.06 [1.04-1.09] | BC | [6] |
| rs2290203 | 15q26.1 | 90968837 | A/G | A | A | *PRC1, PRC1-AS1* | intron variant | 0.504 | $4x10^{-08}$ | 1.08 [1.05-1.11] | BC | [7] |
| rs4784223 | 16q12.1 | 52541995 | A/G | A | G | *TOX3* | intron variant | 0.290 | $6x10^{-21}$ | 1.27 [1.21-1.34] | BC (early onset) | [13] |
| rs3803662 | 16q12.1 | 52552429 | G/A | G | A | *CASC16* | non-coding transcript exon variant | 0.522 | $3x10^{-11}$ | 1.21 [1.15-1.28] | BC | [25] |
| | | | | | | | | 0.260 | $2x10^{-114}$ | 1.24 [1.21-1.27] | BC | [3] |
| | | | | | | | | 0.270 | $1x10^{-09}$ | 1.16 [1.07-1.27] (Het) | BC | [25] |
| | | | | | | | | 0.270 | $6x10^{-19}$ | 1.28 [1.21-1.35] | BC | [9] |
| | | | | | | | | 0.250 | $1x10^{-36}$ | 1.20 [1.16-1.24] | BC | [16] |
| | | | | | | | | 0.267 | $6x10^{-13}$ | 1.14 [1.10-1.18] | BC | [2] |
| | | | | | | | | 0.260 | $3x10^{-15}$ | 1.30 [1.22-1.39] | BC | [12] |
| | | | | | | | | — | $4x10^{-15}$ | 1.50 [1.35-1.66] | BC (male) | [30] |
| rs4784227 | 16q12.1 | 52565276 | C/T | C | T | *CASC16* | intron variant | 0.310 | $3x10^{-09}$ | 1.38 [1.24-1.54] | BC | [21] |
| | | | | | | | | 0.240 | $1x10^{-28}$ | 1.24 [1.20-1.29] | BC | [31] |
| rs12922061 | 16q12.2 | 52601088 | C/T | C | T | *CASC16* | intron variant | 0.241 | $4x10^{-10}$ | 1.23 [1.15-1.31] | BC | [25] |
| rs3112612 | 16q12.2 | 52601252 | G/A | A | G | *CASC16* | intron variant | 0.430 | $4x10^{-10}$ | 1.15 [1.10-1.21] | BC | [11] |
| rs17817449 | 16q12.2 | 53779455 | G/T | G | G | *FTO* | intron variant | 0.600 | $6x10^{-14}$ | 1.08 [1.05-1.10] | BC | [3] |
| rs11075995 | 16q12.2 | 53821379 | T/A | T | A | *FTO* | intron variant | 0.240 | $4x10^{-08}$ | 1.11 [1.07-1.15] | BC | [2] |
| rs13329835 | 16q23.2 | 80616908 | A/G | G | G | *CDYL2* | intron variant | 0.220 | $2x10^{-16}$ | 1.08 [1.05-1.10] | BC | [3] |
| rs146699004* | 17q11.2 | 29230520 | GGT/G | — | — | No mapped genes | | 0.80 | $3x10^{-8}$ | 1.08 [1.04-1.10] | BC | [6] |
| rs12601991 | 17q12 | 37741642 | G/T | G | T | *HNF1B* | intron variant | — | $1x10^{-09}$ | 1.10 [ — ] | Cancer | [1] |
| | | | | | | | | — | $7x10^{-28}$ | — | Cancer (pleiotropy) | [1] |
| rs6504950 | 17q22 | 54979110 | A/G | A | A | *STXBP4* | intron variant | 0.720 | $2x10^{-13}$ | 1.06 [1.04-1.09] | BC | [3] |
| rs745570* | 17q25.3 | 79807926 | G/A | G | A | *CBX8 - LOC102723961* | intergenic variant | 0.500 | $1x10^{-09}$ | 1.05 [1.03-1.08] | BC | [6] |
| rs527616 | 18q11.2 | 26757460 | C/G | G | C | *PCAT18 - LOC105372035* | intron variant | 0.620 | $2x10^{-10}$ | 1.05 [1.03-1.08] | BC | [3] |
| rs1436904 | 18q11.2 | 26990703 | G/T | T | G | *CHST9* | intron variant | 0.600 | $3x10^{-08}$ | 1.04 [1.02-1.06] | BC | [3] |
| rs6507583* | 18q12.3 | 44819625 | G/A | G | G | *SETBP1* | intron variant | 0.930 | $3x10^{-08}$ | 1.10 [1.05-1.14] | BC | [6] |
| rs8170 | 19p13.11 | 17278895 | G/A | G | A | *BABAM1* | synonymous variant | 0.170 | $2x10^{-09}$ | 1.26 [1.17-1.35] | BC | [32] |
| | | | | | | | | 0.191 | $9x10^{-13}$ | 1.15 [1.11-1.20] | BC | [2] |
| | | | A/G | | | | | 0.480 | $4x10^{-13}$ | 1.19 [1.14-1.25] | BC in BRCA1 mutation carriers | [20] |
| rs4808075 | 19p13.11 | 17279482 | T/C | C | C | *BABAM1 - ANKLE1* | intron variant | — | $2x10^{-09}$ | — | Cancer (pleiotropy) | [1] |
| rs8100241 | 19p13.11 | 17282085 | A/G | G | A | *ANKLE1* | missense variant | — | $4x10^{-08}$ | 1.14 [ — ] | BC | [22] |
| rs2363956 | 19p13.11 | 17283315 | T/G | T | G | *ANKLE1* | missense variant | | $2x10^{-08}$ | 1.22 [1.14-1.30] | BC (ER-, PR, and HER-) | [29] |
| rs4808801 | 19p13.11 | 18460331 | G/A | A | G | *ELL* | intron variant | 0.650 | $5x10^{-15}$ | 1.08 [1.05-1.10] | BC | [3] |
| rs3760982 | 19q13.31 | 43782361 | G/A | G | A | *KCNN4 - LOC107987268* | upstream gene variant | 0.460 | $2x10^{-10}$ | 1.06 [1.04-1.08] | BC | [3] |
| rs2284378 | 20q11.22 | 34000289 | C/T | C | T | *RALY* | intron variant | 0.310 | $1x10^{-08}$ | 1.16 [1.10-1.22] | BC | [22] |
| rs2300206 | 20q11.22 | 34002002 | C/G | C | G | *RALY* | intron variant | — | $2x10^{-08}$ | 1.11 [ — ] | Cancer | [1] |
| rs11907546 | 20q11.22 | 34131991 | A/T/C | C | C | *RPS2P1 - ASIP* | regulatory region variant | — | $3x10^{-09}$ | 1.11 [ — ] | Cancer | [1] |
| rs2823093 | 21q21.1 | 15148511 | A/G | A | A | *LOC107985483 - LOC105372739* | intergenic variant | 0.730 | $7x10^{-16}$ | 1.09 [1.06-1.11] | BC | [3] |
| rs16992204 | 21q22.12 | 34738904 | T/C | C | C | *LOC107985515* | upstream gene variant | 0.120 | $5x10^{-08}$ | 1.13 [1.07-1.18] | BC | [4] |
| rs17879961*† | 22q12.1 | 28725099 | A/C/G | A | G | *CHEK2* | missense variant | 0.005 | $1x10^{-8}$ | 1.26 [1.11-1.42] | BC | [6,23] |
| rs132390 | 22q12.2 | 29225488 | T/C | T | C | *EMID1* | intron variant | 0.036 | $3x10^{-09}$ | 1.12 [1.07-1.18] | BC | [3] |
| rs6001930 | 22q13.1 | 40480230 | T/C | C | C | *MKL1* | intron variant | 0.110 | $9x10^{-19}$ | 1.12 [1.09-1.16] | BC | [3] |

* Retrieved from Michailidou et al., 2015
† SNP did not reach genome-wide significance threshold at Michailidou et al., 2015 but was later associated with risk (reported in table; $P \leq 5x10^{-8}$)
[1] dbSNP Build 150
[2] GRCh38
[3] Protective/risk allele, based on the forward strand (Alleles were retrieved from the Ensembl database v91; risk allele is as reported at the GWAS Catalog)
[4] Ancestral allele, retrieved from the Ensembl database v92
[5] Minor allele, retrieved from the Ensembl database v92
[6] Risk allele frequency, reported at the GWAS Catalog
[7] Odds Ratio [95% Confidence Interval], reported at the GWAS Catalog
[8] Trait reported at the GWAS Catalog

**Table S2.** Breast cancer risk associated SNPs located near miRNA genes.

**Table S2a.** Proxy SNPs ($r^2 \geq 0.8$) located downstream or upstream miRNA genes.

| SNP | Locus | Position[1] | Alleles[2] | AA[3] | MA[4] | Gene | Context | GWAS SNP | Distance | $r^2$ | D' |
|---|---|---|---|---|---|---|---|---|---|---|---|
| rs2004659 | 1q21.1 | 145846884 | A/G | A | G | *MIR6736* | downstream gene variant | rs12405132* | 56800 | 0.962 | 1.000 |
| rs9286836 | 1q21.1 | 145846532 | A/C/G | G | G | *MIR6736* | downstream gene variant | rs12405132* | 56448 | 0.926 | 1.000 |
| rs56041970 | 2q34 | 212423018 | T/C | T | C | *MIR548F2* | downstream gene variant | rs13393577 | 9121 | 1.000 | 1.000 |
| rs13413335 | 2q34 | 212423940 | A/C | A | C | *MIR548F2* | downstream gene variant | rs13393577 | 8199 | 0.850 | 1.000 |
| rs10197215 | 2q34 | 212425362 | A/G | A | G | *MIR548F2* | downstream gene variant | rs13393577 | 6777 | 1.000 | 1.000 |
| rs10166403 | 2q34 | 212426475 | A/G | A | G | *MIR548F2* | upstream gene variant | rs13393577 | 5664 | 0.810 | 1.000 |
| rs10202760 | 2q34 | 212426630 | T/C | T | C | *MIR548F2* | upstream gene variant | rs13393577 | 5509 | 1.000 | 1.000 |
| rs77257332 | 2q34 | 212430962 | A/G | A | G | *MIR548F2* | upstream gene variant | rs13393577 | 1177 | 1.000 | 1.000 |
| rs34344935 | 7q21.2 | 92206122 | G/A | G | A | *MIR1285-1* | upstream gene variant | rs6964587*† | 204816 | 0.967 | 1.000 |
| rs1121948 | 8q24.31 | 128152810 | A/G | A | G | *MIR1208* | downstream gene variant | rs11780156 | 29585 | 0.843 | 1.000 |
| rs72843959 | 11p15.5 | 1885061 | C/G | C | G | *MIR7847* | downstream gene variant | rs3817198 | 2715 | 0.964 | 1.000 |
| rs4911146 | 20q11.32 | 34052241 | C/G | G | C | *MIR4755* | downstream gene variant | rs2284378 | 51952 | 1.000 | 1.000 |
| rs3787230 | 20q11.32 | 34052466 | A/G | A | A | *MIR4755* | downstream gene variant | rs2284378 | 52177 | 0.961 | 1.000 |

**Table S2b.** Proxy SNPs ($0.2 \leq r^2 < 0.8$) located downstream or upstream miRNA genes.

| SNP | Locus | Position[1] | Alleles[2] | AA[3] | MA[4] | Gene | Context | GWAS SNP | Distance | $r^2$ | D' |
|---|---|---|---|---|---|---|---|---|---|---|---|
| rs35766535 | 1p36.22 | 10224601 | T/C | T | C | *MIR1273D* | upstream gene variant | rs616488 | 281556 | 0.244 | 0.502 |
| rs61775887 | 1p36.22 | 10224675 | T/C | C | C | *MIR1273D* | upstream gene variant | rs616488 | 281482 | 0.285 | 0.573 |
| rs12122468 | 1p36.22 | 10228364 | A/T | T | T | *MIR1273D* | downstream gene variant | rs616488 | 277793 | 0.285 | 0.573 |
| rs55676616 | 1p36.22 | 10230068 | T/C | C | C | *MIR1273D* | downstream gene variant | rs616488 | 276089 | 0.285 | 0.573 |
| rs17396382 | 1p36.22 | 10230210 | T/C | T | C | *MIR1273D* | downstream gene variant | rs616488 | 275947 | 0.244 | 0.502 |
| rs17400510 | 1p36.22 | 10230363 | T/C | T | C | *MIR1273D* | downstream gene variant | rs616488 | 275794 | 0.261 | 0.530 |
| rs12129035 | 1p36.22 | 10231108 | G/A | G | A | *MIR1273D* | downstream gene variant | rs616488 | 275049 | 0.244 | 0.502 |
| rs12751375 | 1p36.22 | 10231816 | C/G | C | G | *MIR1273D* | downstream gene variant | rs616488 | 274341 | 0.271 | 0.521 |
| rs11133727 | 5p15.33 | 1306650 | C/G | C | – | *MIR4457* | downstream gene variant | rs2736108* | 9277 | 0.246 | 0.682 |
| rs61574973 | 5p15.33 | 1309053 | T/C | C | T | *MIR4457* | downstream gene variant | rs2736108* | 11680 | 0.225 | 0.700 |
| rs6554758 | 5p15.33 | 1310037 | G/A | G | G | *MIR4457* | upstream gene variant | rs2736108* | 12664 | 0.282 | 0.744 |
| rs6866294 | 5p15.33 | 1311578 | T/C | C | T | *MIR4457* | upstream gene variant | rs2736108* | 14205 | 0.264 | 0.695 |
| rs13356727 | 5p15.33 | 1312342 | G/A | G | G | *MIR4457* | upstream gene variant | rs2736108* | 14969 | 0.259 | 0.689 |
| rs759649 | 8q24.21 | 128146998 | G/A | A | G | *MIR1208* | upstream gene variant | rs11780156 | 35397 | 0.214 | 1.000 |
| rs7814495 | 8q24.21 | 128149207 | G/C | G | C | *MIR1208* | upstream gene variant | rs11780156 | 33188 | 0.660 | 1.000 |
| rs10956412 | 8q24.21 | 128150251 | A/C/G | A | C | *MIR1208* | downstream gene variant | rs11780156 | 32144 | 0.598 | 0.851 |
| rs12676304 | 8q24.21 | 128151049 | A/C | A | C | *MIR1208* | downstream gene variant | rs11780156 | 31346 | 0.550 | 1.000 |
| rs759651 | 8q24.21 | 128152042 | T/A/C | T | C | *MIR1208* | downstream gene variant | rs11780156 | 30353 | 0.550 | 1.000 |
| rs1121946 | 8q24.21 | 128152952 | T/G | T | G | *MIR1208* | downstream gene variant | rs11780156 | 29443 | 0.714 | 1.000 |
| rs12675643 | 8q24.21 | 128153446 | T/A | T | A | *MIR1208* | downstream gene variant | rs11780156 | 28949 | 0.550 | 1.000 |
| rs10956413 | 8q24.21 | 128154939 | C/A | C | A | *MIR1208* | downstream gene variant | rs11780156 | 27456 | 0.527 | 1.000 |
| rs5011832 | 10p12.31 | 21493913 | T/C | C | C | *MIR1915* | downstream gene variant | rs7072776 | 250100 | 0.623 | 0.905 |
| rs12770228 | 10p12.31 | 21494705 | G/A | G | A | *MIR1915* | downstream gene variant | rs7072776 | 249308 | 0.697 | 0.869 |
| rs35106872 | 10p12.31 | 21494823 | A/G | G | G | *MIR1915* | downstream gene variant | rs7072776 | 249190 | 0.613 | 0.853 |
| rs12357321 | 10p12.31 | 21501547 | G/A | G | A | *MIR1915* | upstream gene variant | rs7072776 | 242466 | 0.687 | 0.864 |
| rs10839819 | 11p15.5 | 1854956 | C/T | C | T | *MIR4298* | downstream gene variant | rs3817198 | 32820 | 0.227 | 0.900 |
| rs11041481 | 11p15.5 | 1855324 | A/G | G | G | *MIR4298* | downstream gene variant | rs3817198 | 32452 | 0.227 | 0.900 |
| rs10839821 | 11p15.5 | 1855365 | G/A | A | A | *MIR4298* | downstream gene variant | rs3817198 | 32411 | 0.227 | 0.900 |
| rs7934551 | 11p15.5 | 1855487 | C/T | C | T | *MIR4298* | downstream gene variant | rs3817198 | 32289 | 0.227 | 0.900 |
| rs10769814 | 11p15.5 | 1855725 | A/C | C | A | *MIR4298* | downstream gene variant | rs3817198 | 32051 | 0.272 | 0.574 |

| | | | | | | | | | | | |
|---|---|---|---|---|---|---|---|---|---|---|---|
| rs7112859 | 11p15.5 | 1855841 | C/T | C | T | *MIR4298* | downstream gene variant | rs3817198 | 31935 | 0.227 | 0.900 |
| rs869227 | 11p15.5 | 1856208 | A/G | G | G | *MIR4298* | downstream gene variant | rs3817198 | 31568 | 0.227 | 0.900 |
| rs2001487 | 11p15.5 | 1856282 | C/G | G | G | *MIR4298* | downstream gene variant | rs3817198 | 31494 | 0.227 | 0.900 |
| rs907614 | 11p15.5 | 1856303 | T/C | T | T | *MIR4298* | downstream gene variant | rs3817198 | 31473 | 0.272 | 0.574 |
| rs603073 | 11p15.5 | 1856617 | T/C | C | C | *MIR4298* | downstream gene variant | rs3817198 | 31159 | 0.216 | 0.896 |
| rs2685284 | 11p15.5 | 1856843 | G/A | G | A | *MIR4298* | downstream gene variant | rs3817198 | 30933 | 0.227 | 0.900 |
| rs72843933 | 11p15.5 | 1857176 | C/T | C | T | *MIR4298* | downstream gene variant | rs3817198 | 30600 | 0.220 | 0.561 |
| rs1717769 | 11p15.5 | 1857232 | T/C | C | C | *MIR4298* | downstream gene variant | rs3817198 | 30544 | 0.227 | 0.900 |
| rs474016 | 11p15.5 | 1857344 | A/G | G | G | *MIR4298* | downstream gene variant | rs3817198 | 30432 | 0.227 | 0.900 |
| rs599774 | 11p15.5 | 1857406 | A/G | G | G | *MIR4298* | downstream gene variant | rs3817198 | 30370 | 0.227 | 0.900 |
| rs567602 | 11p15.5 | 1859246 | T/C | C | T | *MIR4298* | downstream gene variant | rs3817198 | 28530 | 0.231 | 0.509 |
| rs571122 | 11p15.5 | 1859581 | G/A | A | G | *MIR4298* | upstream gene variant | rs3817198 | 28195 | 0.249 | 0.538 |
| rs587074 | 11p15.5 | 1859841 | G/A | G | G | *MIR4298* | upstream gene variant | rs3817198 | 27935 | 0.206 | 0.471 |
| rs587961 | 11p15.5 | 1860026 | T/C | C | T | *MIR4298* | upstream gene variant | rs3817198 | 27750 | 0.201 | 0.503 |
| rs2048540 | 11p15.5 | 1861018 | T/C | T | T | *MIR4298* | upstream gene variant | rs3817198 | 26758 | 0.249 | 0.538 |
| rs4980392 | 11p15.5 | 1862625 | T/C | T | T | *MIR4298* | upstream gene variant | rs3817198 | 25151 | 0.272 | 0.574 |
| rs72843938 | 11p15.5 | 1863123 | G/A | G | A | *MIR4298* | upstream gene variant | rs3817198 | 24653 | 0.372 | 0.856 |
| rs4980383 | 11p15.5 | 1880867 | C/T | C | T | *MIR7847* | downstream gene variant | rs3817198 | 6909 | 0.385 | 0.876 |
| rs620315 | 11p15.5 | 1881245 | G/A | G | A | *MIR7847* | downstream gene variant | rs3817198 | 6531 | 0.208 | 0.894 |
| rs621679 | 11p15.5 | 1902768 | G/A | G | A | *MIR7847* | downstream gene variant | rs3817198 | 6238 | 0.208 | 0.894 |
| rs61868798 | 11p15.5 | 1883201 | G/A | G | A | *MIR7847* | downstream gene variant | rs3817198 | 4575 | 0.231 | 0.529 |
| rs661348 | 11p15.5 | 1884062 | T/C | T | C | *MIR7847* | downstream gene variant | rs3817198 | 3714 | 0.294 | 0.922 |
| rs3817197 | 11p15.5 | 1884944 | G/A | A | G | *MIR7847* | downstream gene variant | rs3817198 | 2832 | 0.397 | 1.000 |
| rs2009453 | 11q13.1 | 65632057 | C/T | C | T | *MIR4690* | upstream gene variant | rs3903072 | 183538 | 0.299 | 0.680 |
| rs6591183 | 11q13.1 | 65633417 | A/G | G | G | *MIR4690* | upstream gene variant | rs3903072 | 182178 | 0.264 | 0.629 |
| rs10896026 | 11q13.1 | 65633948 | C/T | C | T | *MIR4690* | upstream gene variant | rs3903072 | 181647 | 0.247 | 0.562 |
| rs931127 | 11q13.1 | 65637829 | G/A | G | A | *MIR4690* | downstream gene variant | rs3903072 | 177766 | 0.299 | 0.680 |
| rs2306362 | 11q13.1 | 65638039 | G/A/T | G | T | *MIR4690* | downstream gene variant | rs3903072 | 177556 | 0.247 | 0.562 |
| rs2306364 | 11q13.1 | 65644996 | G/A/C/T | G | A | *MIR4489* | upstream gene variant | rs3903072 | 170599 | 0.350 | 0.602 |
| rs746429 | 11q13.1 | 65649963 | G/A | G | A | *MIR4489* | downstream gene variant | rs3903072 | 165632 | 0.247 | 0.562 |
| rs1466462 | 11q13.1 | 65651893 | G/C | G | C | *MIR4489* | downstream gene variant | rs3903072 | 163702 | 0.226 | 0.510 |
| rs67649296 | 18q11.2 | 26591463 | C/A | C | A | *MIR8057* | downstream gene variant | rs527616 | 165997 | 0.215 | 0.559 |
| rs6059856 | 20q11.22 | 34470149 | G/C | C | C | *MIR644A* | downstream gene variant | rs2284378 | 469859 | 0.413 | 0.933 |
| rs6087587 | 20q11.22 | 34470823 | G/T | G | T | *MIR644A* | downstream gene variant | rs2284378 | 470533 | 0.413 | 0.933 |
| rs79035401 | 22q13.1 | 40816545 | T/C | T | C | *MIR4766* | upstream gene variant | rs6001930 | 336315 | 0.379 | 1.000 |

* Retrieved from Michailidou et al., 2015

† SNP did not reach genome-wide significance threshold at Michailidou et al., 2015 but was later associated with risk ($P \leq 5x10^{-8}$).

[1] GRCh38

[2] Retrieved from the Ensembl database v92

[3] Ancestral allele, retrieved from the Ensembl database v92

[4] Minor allele, retrieved from the Ensembl database v92

**Table S3. Comparison of miRNA-target prediction algorithms.** Existing data or predictions are indicated by a cross.

| Algorithm | Version (Release Year) | Method | | | | Search | Availability | | Ref. |
|---|---|---|---|---|---|---|---|---|---|
| | | Seed Complementarity | Thermodynamics | Conservation | Genomic Context | | Online | Source code or Software | |

| | | | | | | | | | |
|---|---|---|---|---|---|---|---|---|---|
| **TargetScan** | 7.1 (2015) | x | x | x | x | 3'UTR | www.targetscan.org | x (Perl) | [33] |
| **miRanda** | 3.3a (2010) | | x | | | 5'UTR, CDS, 3'UTR | www.microrna.org | x | [34,35] |
| **PicTar** | – (2007) | | x | x | | 3'UTR | pictar.mdc-berlin.de | | [36] |
| **DIANA microT-CDS** | 5.0 (2012) | x | x | x | x | CDS, 3'UTR | www.microrna.gr/microT-CDS | | [37,38] |
| **RNAHybrid** | – (2004) | | x | | | 3'UTR | bibiserv2.cebitec.uni-bielefeld.de/rnahybrid | x | [39,40] |
| **PITA** | 6 (2008) | x | x | x | | 3'UTR | genie.weizmann.ac.il/pubs/mir07 | x (Perl) | [41] |
| **miRTar** | | x | x | x | x | 5'UTR, CDS, 3'UTR | mirtar.mbc.nctu.edu.tw/human | | [42] |

**Table S4. 3'UTR-located BC risk-associated variants.** Existing data or predictions are indicated by a cross.

| Locus | GWAS SNP | LD[1] | | SNP | Alleles | A.A.[2] | M.A.[3] | Gene ID | Gene | Transcript ID | TargetScan v7.1 | miRanda 3.3a |
|---|---|---|---|---|---|---|---|---|---|---|---|---|
| | | $r^2$ | D' | | | | | | | | | |
| 1q21.1 | rs12405132 | 0.851 | 0.959 | rs17354678 | T/C | T | C | ENSG00000265491 | *RNF115* | ENST00000582693 | x | x |
| | | 0.851 | 0.959 | rs12123298 | G/A/C | G | A | | | | x | x |
| | | 0.961 | 1 | rs17352469 | T/C | T | C | | | | x | x |
| | | 0.961 | 1 | rs2231375 | C/T | C | T | ENSG00000117281 | *CD160* | ENST00000616463 | x | x |
| | | 0.961 | 1 | rs1778523 | G/C | G | C | | | | x | x |
| 1q32.1 | rs4245739 | 0.821 | 0.959 | rs4245738 | C/T | T | C | ENSG00000198625 | *MDM4* | ENST00000367180 | x | x |
| | | 1 | 1 | rs4245739 | C/A | A | C | ENSG00000198625 | *MDM4* | ENST00000391947 | x | x |
| | | 1 | 1 | rs4245739 | C/A | A | C | ENSG00000198625 | *MDM4* | ENST00000621032 | x | x |
| | | 1 | 1 | rs4245739 | C/A | A | C | ENSG00000198625 | *MDM4* | ENST00000612738 | x | x |
| | | 1 | 1 | rs4245739 | C/A | A | C | ENSG00000198625 | *MDM4* | ENST00000616250 | x | x |
| | | 1 | 1 | rs4245739 | C/A | A | C | ENSG00000198625 | *MDM4* | ENST00000454264 | x | x |
| | | 1 | 1 | rs4245739 | C/A | A | C | ENSG00000198625 | *MDM4* | ENST00000367182 | x | x |
| | | 1 | 1 | rs4245739 | C/A | A | C | ENSG00000198625 | *MDM4* | ENST00000367183 | x | x |
| | | 1 | 1 | rs4245739 | C/A | A | C | ENSG00000198625 | *MDM4* | ENST00000614459 | x | x |
| | | 0.861 | 1 | rs10900596 | T/C | C | T | ENSG00000198625 | *MDM4* | ENST00000612738 | x | x |
| | | 0.861 | 1 | rs10900596 | T/C | C | T | ENSG00000198625 | *MDM4* | ENST00000367183 | x | x |
| | | 0.861 | 1 | rs10900596 | T/C | C | T | ENSG00000198625 | *MDM4* | ENST00000367182 | x | x |
| | | 0.861 | 1 | rs10900596 | T/C | C | T | ENSG00000198625 | *MDM4* | ENST00000391947 | x | x |
| | | 0.861 | 1 | rs10900596 | T/C | C | T | ENSG00000198625 | *MDM4* | ENST00000614459 | x | x |
| | | 0.861 | 1 | rs10900596 | T/C | C | T | ENSG00000198625 | *MDM4* | ENST00000616250 | x | x |
| | | 0.861 | 1 | rs10900596 | T/C | C | T | ENSG00000198625 | *MDM4* | ENST00000454264 | x | x |
| | | 0.861 | 1 | rs10900596 | T/C | C | T | ENSG00000198625 | *MDM4* | ENST00000621032 | x | x |
| | | 0.861 | 1 | rs10900597 | C/T | C | C | ENSG00000198625 | *MDM4* | ENST00000621032 | x | x |
| | | 0.861 | 1 | rs10900597 | C/T | C | C | ENSG00000198625 | *MDM4* | ENST00000367183 | x | x |
| | | 0.861 | 1 | rs10900597 | C/T | C | C | ENSG00000198625 | *MDM4* | ENST00000454264 | x | x |
| | | 0.861 | 1 | rs10900597 | C/T | C | C | ENSG00000198625 | *MDM4* | ENST00000614459 | x | x |
| | | 0.861 | 1 | rs10900597 | C/T | C | C | ENSG00000198625 | *MDM4* | ENST00000367182 | x | x |
| | | 0.861 | 1 | rs10900597 | C/T | C | C | ENSG00000198625 | *MDM4* | ENST00000391947 | x | x |
| | | 0.861 | 1 | rs10900597 | C/T | C | C | ENSG00000198625 | *MDM4* | ENST00000616250 | x | x |
| | | 0.861 | 1 | rs10900597 | C/T | C | C | ENSG00000198625 | *MDM4* | ENST00000612738 | x | x |
| 3p24.1 | rs4973768 | 1 | 1 | rs4973768 | C/T | T | T | ENSG00000033867 | *SLC4A7* | ENST00000428386 | x | x |
| | | 1 | 1 | rs4973768 | C/T | T | T | ENSG00000033867 | *SLC4A7* | ENST00000295736 | x | x |
| | | 1 | 1 | rs4973768 | C/T | T | T | ENSG00000033867 | *SLC4A7* | ENST00000419036 | x | x |
| | | 1 | 1 | rs4973768 | C/T | T | T | ENSG00000033867 | *SLC4A7* | ENST00000425128 | x | x |
| | rs653465 | 0.839 | 0.963 | rs4973768 | C/T | T | T | ENSG00000033867 | *SLC4A7* | ENST00000419036 | x | x |
| | | 0.839 | 0.963 | rs4973768 | C/T | T | T | ENSG00000033867 | *SLC4A7* | ENST00000295736 | x | x |

| | | | | | | | | | | | | |
|---|---|---|---|---|---|---|---|---|---|---|---|---|
| | | 0.839 | 0.963 | rs4973768 | C/T | T | T | ENSG00000033867 | *SLC4A7* | ENST00000428386 | x | x |
| | | 0.839 | 0.963 | rs4973768 | C/T | T | T | ENSG00000033867 | *SLC4A7* | ENST00000425128 | x | x |
| | rs4973768 | 1 | 1 | rs1051545 | T/C | C | C | ENSG00000033867 | *SLC4A7* | ENST00000295736 | x | x |
| | | 1 | 1 | rs1051545 | T/C | C | C | ENSG00000033867 | *SLC4A7* | ENST00000425128 | x | x |
| | | 1 | 1 | rs1051545 | T/C | C | C | ENSG00000033867 | *SLC4A7* | ENST00000419036 | x | x |
| | | 1 | 1 | rs1051545 | T/C | C | C | ENSG00000033867 | *SLC4A7* | ENST00000428386 | x | x |
| | rs653465 | 0.839 | 0.963 | rs1051545 | T/C | C | C | ENSG00000033867 | *SLC4A7* | ENST00000295736 | x | x |
| | | 0.839 | 0.963 | rs1051545 | T/C | C | C | ENSG00000033867 | *SLC4A7* | ENST00000428386 | x | x |
| | | 0.839 | 0.963 | rs1051545 | T/C | C | C | ENSG00000033867 | *SLC4A7* | ENST00000425128 | x | x |
| | | 0.839 | 0.963 | rs1051545 | T/C | C | C | ENSG00000033867 | *SLC4A7* | ENST00000419036 | x | x |
| 3p14.1 | rs1053338 | 0.895 | 1 | rs3733126 | C/T | T | T | ENSG00000285258 | *ATXN7* | ENST00000295900 | x | x |
| | | 0.895 | 1 | rs3733126 | C/T | T | T | ENSG00000163635 | *ATXN7* | ENST00000538065 | x | x |
| | | 0.837 | 0.941 | rs1046025 | C/T | T | T | ENSG00000163636 | *PSMD6* | ENST00000480205 | x | x |
| | | 0.837 | 0.941 | rs1046025 | C/T | T | T | ENSG00000163636 | *PSMD6* | ENST00000482510 | x | x |
| | | 0.837 | 0.941 | rs1046025 | C/T | T | T | ENSG00000163636 | *PSMD6* | ENST00000492933 | x | x |
| | | 0.837 | 0.941 | rs1046025 | C/T | T | T | ENSG00000163636 | *PSMD6* | ENST00000295901 | x | x |
| | | 0.837 | 0.941 | rs1046025 | C/T | T | T | ENSG00000163636 | *PSMD6* | ENST00000394431 | x | x |
| 5q11.2 | rs12655019 | 0.92 | 1 | rs1466010 | A/G | A | G | ENSG00000155542 | *SETD9* | ENST00000418299 | x | x |
| | | 0.92 | 1 | rs12654125 | G/A | G | A | ENSG00000155545 | *MIER3* | ENST00000381226 | x | x |
| | | 0.92 | 1 | rs12654125 | G/A | G | A | ENSG00000155545 | *MIER3* | ENST00000452157 | x | x |
| | | 0.92 | 1 | rs12654125 | G/A | G | A | ENSG00000155545 | *MIER3* | ENST00000381199 | x | x |
| | | 0.92 | 1 | rs12654125 | G/A | G | A | ENSG00000155545 | *MIER3* | ENST00000381213 | x | x |
| | | 0.92 | 1 | rs3756586 | A/G | G | G | ENSG00000155545 | *MIER3* | ENST00000381226 | x | x |
| | | 0.92 | 1 | rs3756586 | A/G | G | G | ENSG00000155545 | *MIER3* | ENST00000381213 | x | x |
| | | 0.92 | 1 | rs3756586 | A/G | G | G | ENSG00000155545 | *MIER3* | ENST00000381199 | x | x |
| | | 0.92 | 1 | rs3756586 | A/G | G | G | ENSG00000155545 | *MIER3* | ENST00000452157 | x | x |
| | | 0.92 | 1 | rs16886496 | T/C | T | C | ENSG00000155545 | *MIER3* | ENST00000381226 | x | x |
| | | 0.92 | 1 | rs16886496 | T/C | T | C | ENSG00000155545 | *MIER3* | ENST00000381199 | x | x |
| | | 0.92 | 1 | rs16886496 | T/C | T | C | ENSG00000155545 | *MIER3* | ENST00000381213 | x | x |
| | | 0.92 | 1 | rs16886496 | T/C | T | C | ENSG00000155545 | *MIER3* | ENST00000452157 | x | x |
| 5q14.2 | rs7707921 | 0.881 | 1 | rs73136782 | T/G | T | G | ENSG00000152348 | *ATG10* | ENST00000355178 | x | x |
| | | 1 | 1 | rs6884232 | G/A | A | G | ENSG00000152348 | *ATG10* | ENST00000282185 | x | x |
| | | 1 | 1 | rs6884232 | G/A | A | G | ENSG00000152348 | *ATG10* | ENST00000458350 | x | x |
| | | 1 | 1 | rs1019806 | G/A | A | G | ENSG00000152348 | *ATG10* | ENST00000282185 | x | x |
| 6q22.33 | rs2180341 | 1 | 1 | rs9321073 | C/T | T | C | ENSG00000118518 | *RNF146* | ENST00000356799 | x | x |
| | | 1 | 1 | rs9321073 | C/T | T | C | ENSG00000118518 | *RNF146* | ENST00000309649 | x | x |
| | | 1 | 1 | rs9321073 | C/T | T | C | ENSG00000118518 | *RNF146* | ENST00000368314 | x | x |
| | | 1 | 1 | rs9321073 | C/T | T | C | ENSG00000118518 | *RNF146* | ENST00000616343 | x | x |
| 6q25.1 | rs12662670 | 0.892 | 1 | rs3734805 | A/C | A | C | ENSG00000120262 | *CCDC170* | ENST00000239374 | x | x |
| | | 0.892 | 1 | rs9383935 | C/T | C | T | ENSG00000120262 | *CCDC170* | ENST00000239374 | x | x |
| | | 0.892 | 1 | rs9383589 | A/G | A | G | ENSG00000120262 | *CCDC170* | ENST00000239374 | x | x |
| | rs2046210 | 0.821 | 1 | rs3734806 | G/A | G | A | ENSG00000120262 | *CCDC170* | ENST00000239374 | x | x |
| | | 0.821 | 1 | rs3757322 | T/G | T | G | ENSG00000120262 | *CCDC170* | ENST00000239374 | x | x |
| 7q21.2 | rs6964587 | 1 | 1 | rs55745934 | T/C | T | C | ENSG00000127914 | *AKAP9* | ENST00000619023[†] | | x |
| | | 1 | 1 | rs10225885 | A/G | A | G | ENSG00000127914 | *AKAP9* | ENST00000619023[†] | | x |
| | | 1 | 1 | rs10225892 | A/G | G | G | ENSG00000127914 | *AKAP9* | ENST00000619023[†] | | x |
| | | 1 | 1 | rs28584017 | G/A | G | A | ENSG00000127914 | *AKAP9* | ENST00000435423[†] | | x |
| | | 1 | 1 | rs28584017 | G/A | G | A | ENSG00000127914 | *AKAP9* | ENST00000358100[†] | | x |
| | | 1 | 1 | rs4265 | C/T | C | T | ENSG00000127914 | *AKAP9* | ENST00000358100[†] | | x |
| | | 1 | 1 | rs4265 | C/T | C | T | ENSG00000127914 | *AKAP9* | ENST00000435423[†] | | x |

| | | | | | | | | | | | | |
|---|---|---|---|---|---|---|---|---|---|---|---|---|
| 11q13.1 | rs3903072 | 0.837 | 0.963 | rs633800 | G/A | G | A | ENSG00000172638 | *EFEMP2* | ENST00000530850* | | x |
| | | 0.837 | 0.963 | rs633800 | G/A | G | A | ENSG00000172638 | *EFEMP2* | ENST00000533347* | | x |
| 13q13.1 | rs56084662 | – | – | rs56084662 | G/A | G | A | ENSG00000073910 | *FRY* | ENST00000380250 | x | x |
| | | – | – | rs56084662 | G/A | G | A | ENSG00000073910 | *FRY* | ENST00000645780 | x | x |
| | | – | – | rs56084662 | G/A | G | A | ENSG00000073910 | *FRY* | ENST00000542859 | x | x |
| | | – | – | rs56084662 | G/A | G | A | ENSG00000073910 | *FRY* | ENST00000642040 | x | x |
| | | – | – | rs56084662 | G/A | G | A | ENSG00000073910 | *FRY* | ENST00000647500 | x | X |
| 15q26.1 | rs2290203 | 1 | 1 | rs2290203 | G/A | A | A | ENSG00000284946 | – | ENST00000643536* | | x |
| | | 0.938 | 1 | rs2301826 | C/T | T | T | ENSG00000284946 | – | ENST00000647331* | | x |
| | | 0.938 | 1 | rs2301826 | C/T | T | T | ENSG00000284946 | – | ENST00000643536* | | x |
| 17q22 | rs6504950 | 0.83 | 1 | rs3087650 | G/A | G | A | ENSG00000166260 | *COX11* | ENST00000576370* | | x |
| | | 0.83 | 1 | rs3087650 | G/A | G | A | ENSG00000166260 | *COX11* | ENST00000574821* | | x |
| | | 0.83 | 1 | rs3087650 | G/A | G | A | ENSG00000166260 | *COX11* | ENST00000572558* | | x |
| | | 0.83 | 1 | rs1802212 | A/C | A | C | ENSG00000141198 | *TOM1L1* | ENST00000575882 | x | x |
| | | 0.83 | 1 | rs1802212 | A/C | A | C | ENSG00000141198 | *TOM1L1* | ENST00000445275 | x | x |
| | | 0.83 | 1 | rs1802212 | A/C | A | C | ENSG00000166260 | *COX11* | ENST00000299335 | x | x |
| | | 0.83 | 1 | rs1802212 | A/C | A | C | ENSG00000141198 | *TOM1L1* | ENST00000348161 | x | x |
| | | 0.83 | 1 | rs1802212 | A/C | A | C | ENSG00000141198 | *TOM1L1* | ENST00000536554 | x | x |
| | | 0.83 | 1 | rs1802212 | A/C | A | C | ENSG00000166260 | *COX11* | ENST00000576370 | x | x |
| | | 0.83 | 1 | rs1802212 | A/C | A | C | ENSG00000141198 | *TOM1L1* | ENST00000572158 | x | x |
| | | 0.83 | 1 | rs1802212 | A/C | A | C | ENSG00000141198 | *TOM1L1* | ENST00000571319 | x | x |
| | | 0.83 | 1 | rs17817901 | A/G | A | G | ENSG00000141198 | *TOM1L1* | ENST00000445275 | x | x |
| | | 0.83 | 1 | rs17817901 | A/G | A | G | ENSG00000166260 | *COX11* | ENST00000576370 | x | x |
| | | 0.83 | 1 | rs17817901 | A/G | A | G | ENSG00000166260 | *COX11* | ENST00000299335 | x | x |
| | | 0.83 | 1 | rs17817901 | A/G | A | G | ENSG00000141198 | *TOM1L1* | ENST00000571319 | x | x |
| | | 0.83 | 1 | rs17817901 | A/G | A | G | ENSG00000141198 | *TOM1L1* | ENST00000536554 | x | x |
| | | 0.83 | 1 | rs17817901 | A/G | A | G | ENSG00000141198 | *TOM1L1* | ENST00000348161 | x | x |
| | | 0.83 | 1 | rs17817901 | A/G | A | G | ENSG00000141198 | *TOM1L1* | ENST00000575882 | x | x |
| 19p13.11 | rs8170 | 1 | 1 | rs8170 | G/A | G | A | ENSG00000105393 | *BABAM1* | ENST00000601232* | | x |
| | rs2363956 | 1 | 1 | rs8100241 | G/A | G | A | ENSG00000269307 | – | ENST00000596542* | | x |
| | rs8100241 | 1 | 1 | rs8100241 | G/A | G | A | ENSG00000269307 | – | ENST00000596542* | | x |
| | rs2363956 | 1 | 1 | rs8108174 | T/A | T | A | ENSG00000269307 | – | ENST00000596542* | | x |
| | | 1 | 1 | rs8108174 | T/A | T | A | ENSG00000160117 | *ANKLE1* | ENST00000404085* | | x |
| | rs8100241 | 1 | 1 | rs8108174 | T/A | T | A | ENSG00000269307 | – | ENST00000596542* | | x |
| | rs8100241 | 1 | 1 | rs8108174 | T/A | T | A | ENSG00000160117 | *ANKLE1* | ENST00000404085* | | x |
| | rs2363956 | 1 | 1 | rs2363956 | T/G | T | G | ENSG00000160117 | *ANKLE1* | ENST00000404085* | | x |
| | rs8100241 | 1 | 1 | rs2363956 | T/G | T | G | ENSG00000160117 | *ANKLE1* | ENST00000404085* | | x |
| | rs8170 | 1 | 1 | rs10425939 | C/T | C | T | ENSG00000160117 | *ANKLE1* | ENST00000404085 | x | x |
| | rs8170 | 1 | 1 | rs10425939 | C/T | C | T | ENSG00000160117 | *ANKLE1* | ENST00000404261 | x | x |
| | rs4808075 | 1 | 1 | rs4808616 | C/A | C | A | ENSG00000127220 | *ABHD8* | ENST00000247706 | x | x |
| | rs8170 | 0.859 | 0.95 | rs11540855 | A/G/T | A | G | ENSG00000127220 | *ABHD8* | ENST00000247706 | x | x |
| | rs4808801 | 1 | 1 | rs10405636 | A/C | C | C | ENSG00000130511 | *SSBP4* | ENST00000607020* | | x |
| | | 1 | 1 | rs10405636 | A/C | C | C | ENSG00000130511 | *SSBP4* | ENST00000601614* | | x |
| | | 0.965 | 1 | rs2385088 | A/G | G | G | ENSG00000105655 | *ISYNA1* | ENST00000338128 | x | x |
| | | 0.965 | 1 | rs2385088 | A/G | G | G | ENSG00000105655 | *ISYNA1* | ENST00000582811 | x | x |
| | | 1 | 1 | rs2303697 | T/C | C | C | ENSG00000105655 | *ISYNA1* | ENST00000582770* | | x |
| | | 1 | 1 | rs2303697 | T/C | C | C | ENSG00000105655 | *ISYNA1* | ENST00000582811* | | x |
| | | 1 | 1 | rs2303697 | T/C | C | C | ENSG00000105655 | *ISYNA1* | ENST00000577820* | | x |
| | | 0.897 | 1 | rs1043327 | A/G | G | G | ENSG00000105656 | *ELL* | ENST00000594635 | x | x |
| | | 0.897 | 1 | rs1043327 | A/G | G | G | ENSG00000105656 | *ELL* | ENST00000262809 | x | x |

| | | | | | | | | | | | | |
|---|---|---|---|---|---|---|---|---|---|---|---|---|
| 20q11.22 | rs2284378 | 1 | 1 | rs6119447 | A/G | G | A | ENSG00000125970 | *RALY* | ENST00000375114 | x | x |
| | | 1 | 1 | rs8123521 | A/C | C | A | ENSG00000125970 | *RALY* | ENST00000375114 | x | x |
| 22q12.1 | rs17879961 | – | – | rs17879961 | A/C/G | A | G | ENSG00000183765 | *CHEK2* | ENST00000454252* | | x |

* Non-sense mediated decay transcript (Ensembl v92)
† Transcript not available at TargetScan's dataset of 3'UTRs whole-genome alignments
[1] Linkage disequilibrium, obtained from SNAP (Pilot release of 1000 Genomes Project; CEU population; $r^2 \geq 0.8$; distance limit = 500 kb)
[2] Ancestral allele, retrieved from the Ensembl database v92
[3] Minor allele, retrieved from the Ensembl database v92

**Figure S1. GTEx expression (v7) of transcripts predicted to be affected by candidate rSNPs at 19p13.11 and 1q21.1 loci in breast (mammary tissue).** Panel to the left indicates total expression levels for the gene. Panel to the right indicates expression levels for all transcripts of each gene individually. Expression levels are in Transcripts per Million (TPM).

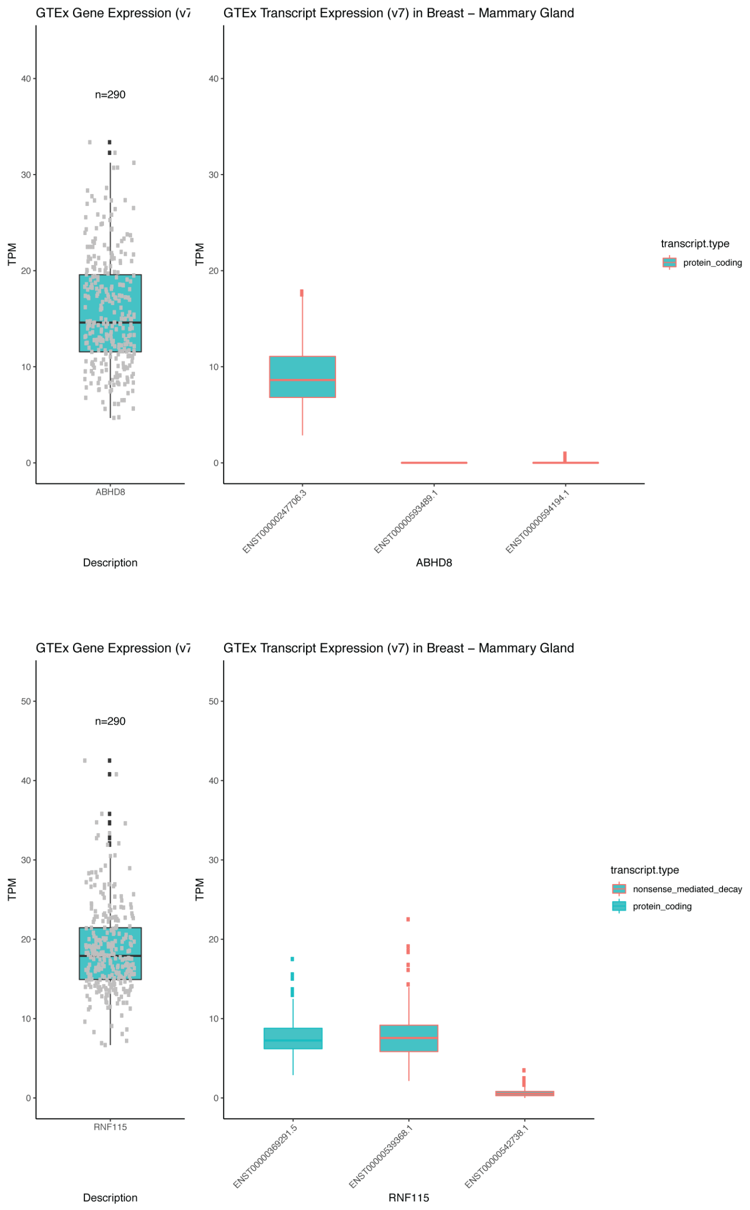


**Figure S2. DAE data for *ATG10*, *CCDC170* and *ISYNA1* in breast (mammary tissue).** DAE data for heterozygous individuals tested for the SNP indicated immediately below. The alleles are indicated for each SNP in the order of the AE ratio calculated (i.e. A/G corresponds to the ratio of allele A by allele G). Dashed horizontal lines indicate the threshold for DAE set at 1.5 fold difference between alleles (|$\log_2$ AE$_{ratio}$|= 0.58).

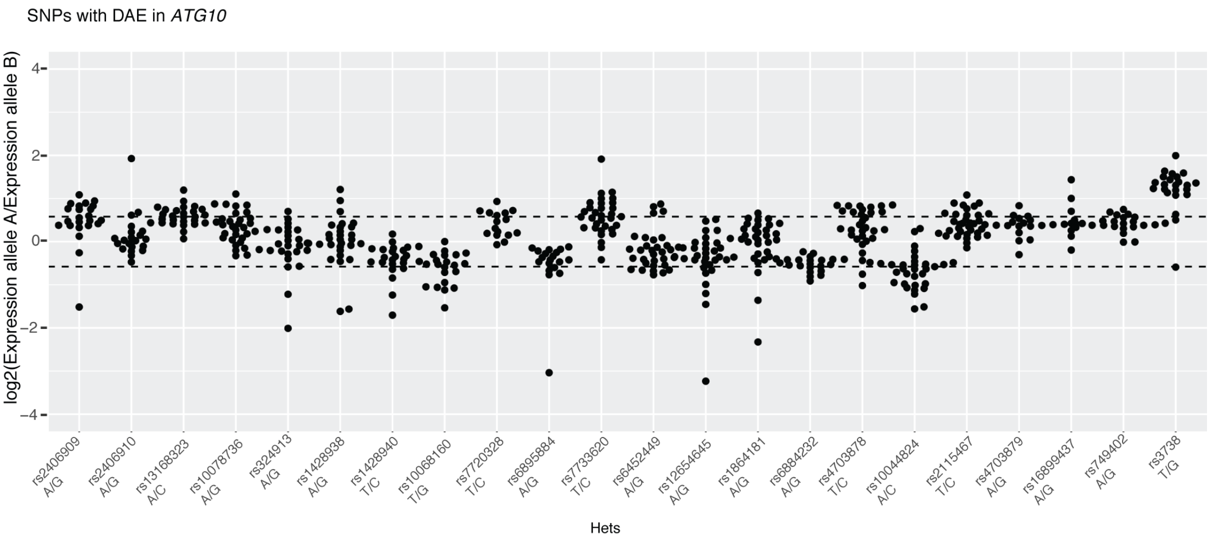


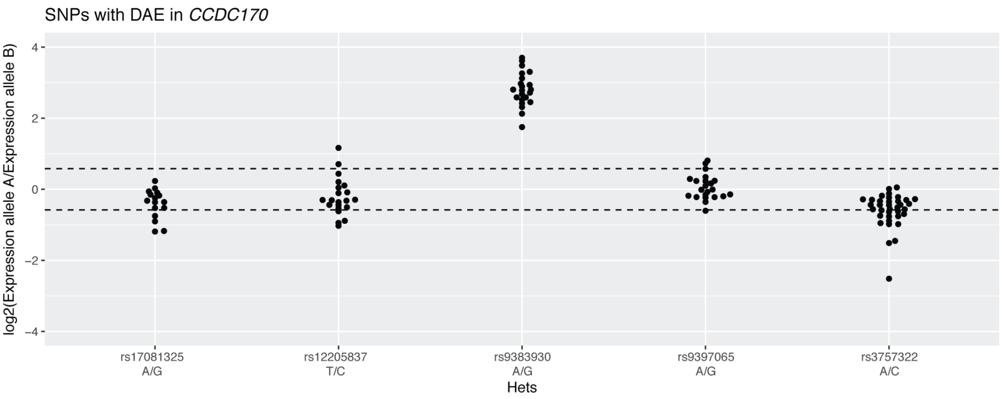


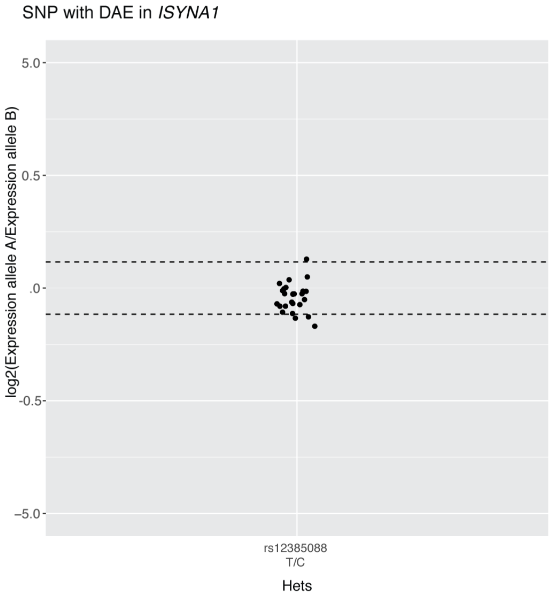

33. Agarwal V, Bell GW, Nam J-W, Bartel DP. Predicting effective microRNA target sites in mammalian

mRNAs. Elife [Internet]. 2015;4:1–38. Available from: http://elifesciences.org/lookup/doi/10.7554/eLife.05005
